# Somatic gene delivery for flexible *in vivo* modeling of high-risk sarcoma

**DOI:** 10.1101/2024.01.30.577924

**Authors:** Roland Imle, Daniel Blösel, Felix K.F. Kommoss, Eric Stutheit Zhao, Robert Autry, Christina Blume, Dmitry Lupar, Lukas Schmitt, Claudia Winter, Lena Wagner, Sara Placke, Malte von Eicke, Michael Hertwig, Heike Peterziel, Ina Oehme, Sophia Scheuerman, Christian Seitz, Florian H. Geyer, Florencia Cidre-Aranaz, Thomas G. P. Grünewald, Christian Vokuhl, Priya Chudasama, Claudia Scholl, Claudia Schmidt, Patrick Günther, Martin Sill, Kevin B. Jones, Stefan M. Pfister, Ana Banito

## Abstract

A particular challenge hampering therapeutic advancements for high-risk sarcoma patients is the broad spectrum of molecularly distinct sarcoma entities and the corresponding lack of suitable model systems to recapitulate and study these diseases. To overcome this predicament, we developed a novel genetically-controlled, yet versatile mouse modeling platform allowing delivery of different genetic lesions by electroporation (EPO) of the thigh muscle wildtype mice. This optimized sarcoma EPO-GEMM (EPO-based genetically engineered mouse model) platform allowed the generation of ten biologically distinct sarcoma entities, including Synovial Sarcoma (SS), fusion-positive and fusion-negative Rhabdomyosarcoma (RMS), Alveolar Soft Part Sarcoma (ASPS), Undifferentiated Pleomorphic Sarcoma (UPS) and Infantile Fibrosarcoma (IFS). Comprehensive molecular profiling and cross-species analyses confirmed faithful recapitulation of the human disease, including the expression of relevant immunotherapy targets. Syngeneic allografting enabled reliable preservation and scalability of Sarcoma-EPO-GEMMs for treatment trials, such as B7-H3-directed CAR-T cell therapy in an immunocompetent background.

## INTRODUCTION

Sarcomas are a group of mesenchymal cancers arising in soft tissues or bone that disproportionately often affect children, adolescents, and young adults (Grünewald et al., 2020). While sarcomas arising from bone mostly fall into the categories of Osteosarcoma (OS), Ewing Sarcoma (EwS) and Chondrosarcoma (CS), Soft-tissue Sarcomas (STS) are significantly more diverse with more than 50 genetically distinct entities and even more subtypes. They are broadly classified into Rhabdomyosarcomas (RMS) which retain some skeletal muscle differentiation and Non-Rhabdomyosarcomas STS (NRSTS) consisting of all remaining STS entities with diverse subtypes (Antonescu et al., 2020; Pfister et al., 2022). Many sarcoma entities display a low tumor mutational burden (TMB) and are often driven by dominant fusion oncoproteins involving chromatin associated regulators and transcription factors (Perry et al., 2019). Others are characterized by chromosomal instability and high TMB. Unlike many other cancers, substantial therapeutic and prognostic advances have largely been lacking for sarcomas for several decades across all age groups. This is a result of various factors including their immense heterogeneity and lack of clinically actionable targets (Sandler et al., 2019). Clinical management of sarcomas essentially relies on extensive empirical experience in multiagent chemotherapy combined with surgery and irradiation. Although few targeted therapies or immunotherapies have proven efficacious against sarcomas (van Tilburg et al., 2021), they exhibit differential responses to cytotoxic agents stemming from unknown entity-specific mechanisms of oncogenesis and progression.

A major bottleneck to achieve long-sought therapeutic advances for sarcoma patients is the lack of adequate model systems for mechanistic and preclinical studies (Jones et al., 2019). Modeling sarcoma is particularly challenging. Due to their rarity and heterogeneity, patient tissue is notoriously scarce, making derivative models poorly available. Genetically- engineered mouse models (GEMMs) are difficult to establish given that the exact cell of origin remains elusive for most entities (Kannan et al., 2021) and widespread expression of many sarcoma drivers results in embryonic lethality (Haldar et al., 2007). Yet, several sarcoma GEMMs could be established by introducing mutations into the germline (Imle et al., 2021), often requiring conditional Cre recombinase expression in defined lineages and developmental windows (Han et al., 2016). Germline GEMMs typically exhibit multifocal tumor onset under mixed genetic backgrounds and have generally been impractical for preclinical testing. Aiming to overcome these challenges in sarcoma modeling, we utilized *in vivo* electroporation (EPO) of CRISPR and transposon vectors as a somatic gene delivery approach (Lima and Maddalo, 2021) to murine soft-tissue in wildtype mice. We envisioned that the speedy and cell-type-agnostic nature of this technique would allow us to model genetically-defined sarcomas of myogenic (RMS) and non-myogenic (NRSTS) origin with localized onset in the physiological context of a fully competent immune system. We applied this optimized method to study various sarcoma-associated genetic alterations and successfully generated a broad range of more than ten fusion-positive and -negative sarcoma mouse models, including the first NTRK-driven GEMM. We also provide novel experimental evidence that *Bcor*, which is inactivated in 15-20% of RMS (Shern et al., 2021), can function as a tumor suppressor in fusion-negative RMS. The array of models recapitulates diverse tumorigenic mechanisms ranging from rewiring of epigenetic machinery by gene fusions (*SS18::SSX1/2*), to uncoupled tyrosine kinase activation (*ETV6::NTRK3*), to aberrant transcription factors (*ASPSCR1::TFE3*), and even multistep tumorigenesis based on *Trp53*- mutation-related genomic instability and second hit mutations (e.g. *Nf1* or *Smarcb1*). After validating the reliability of EPO-GEMMs by multiomic cross-species analysis, we further derived syngeneic allograft models (SAMs) to preserve, share and scale this one-of-a-kind collection of newly established sarcoma GEMMs. We demonstrate the feasibility of preclinical testing in mouse sarcoma cell lines and immunocompetent sarcoma SAMs using NTRK inhibitor treatment in Infantile Fibrosarcoma (IFS), as well as chimeric antigen receptor T lymphocyte (CAR-T) immunotherapy targeting B7-H3 in alveolar RMS (aRMS).

## RESULTS

### An optimized protocol allows efficient *in vivo* genetic manipulation of mouse muscle tissue

To establish an optimized protocol for orthotopic induction of murine STS by electroporation, we chose a vector combination conveying overexpression of oncogenic *RAS* and inactivation of *Trp53*, which have been shown to co-occur in several high-risk sarcoma entities, such as embryonal RMS (eRMS) and have previously been validated to efficiently drive sarcoma in mice from the *Rosa26* locus upon Cre-LoxP recombination (Dodd et al., 2015). We performed survival surgery to expose the thigh muscle in 4–6-week-old mice (referred to as P30 hereafter) and delivered a plasmid mix containing (i) Sleeping Beauty (SB13) transposase, (ii) a transposon vector expressing oncogenic *KRAS* (K), and luciferase/GFP reporter genes (either as an all-in-one vector (KGL) or in separate plasmids (K+L), (iii) a CRISPR/Cas9 vector expressing Cas9 endonuclease and a single-guide RNA (sgRNA) targeting *Trp53* **(Fig. 1A-C)**. The procedure was performed bilaterally, and methylene blue dye was used to demarcate the quadriceps muscles before EPO with two 5 mm plate electrodes. To expand the range of accessible cell types and the developmental window for oncogenic transformation, the EPO procedure was also established in neonatal (P0) animals. Here, plasmid injection and EPO were performed transcutaneously **(Fig. 1A)**. These initial electroporation attempts were unsuccessful in inducing tumors, underscoring the previously noted low efficiency of muscle transfection (Wang et al., 2005) **(Table S1)**. We therefore optimized a dedicated protocol for somatic muscle engineering to induce sarcomagenesis using *in vivo* bioluminescence imaging (IVIS) two days and one week post EPO as a surrogate for transfection efficiency using different EPO conditions **(Fig. S1A)**. Optimal EPO conditions were determined as five unilateral pulses of 100V for 35ms for P30 and 70V for P0 animals. Pre-treatment with hyaluronidase (Hyal) to temporarily loosen up the extracellular matrix (McMahon et al., 2001), as well as delivering oncogenes and reporter genes on separate plasmids (K+L versus KGL) further increased transfection efficiency **(Fig. S1A)**. The additionally tested resuspension of plasmid DNA in cationic polymer Poly-L- glutamate (Glut) (Nicol et al., 2002) did not show added benefit in this setting **(Fig. S1A)**.

**Figure. 1.**
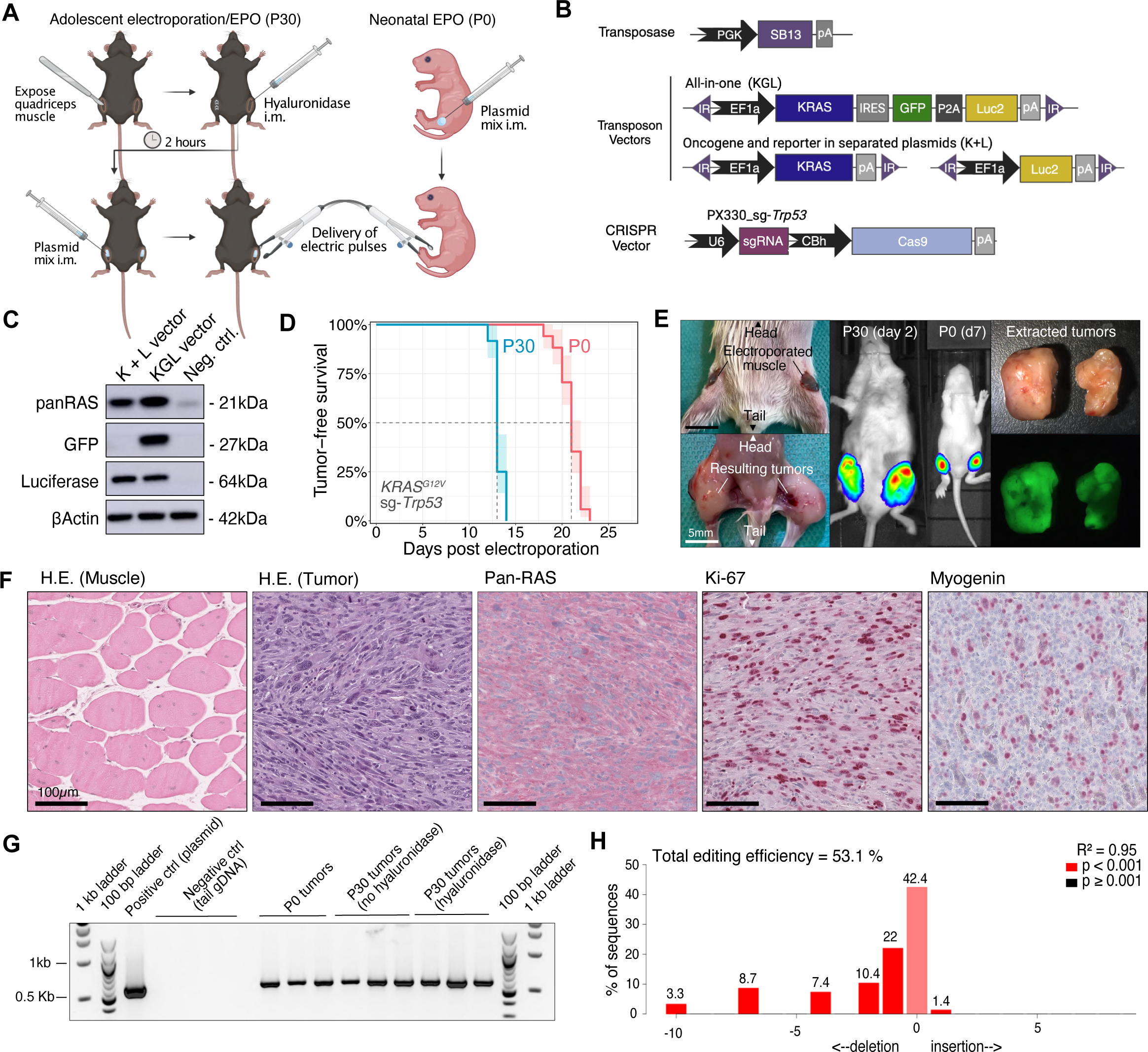
An optimized protocol allows efficient *in vivo* genetic manipulation of mouse muscle tissue A) Scheme of electroporation procedure in P30 and P0 animals. B) Illustration of utilized transposon and CRISPR vectors. KGL = EF1a-KRAS-IRES-eGFP-P2A-Luc; K+L = EF1a- KRAS + PGK-Luc; C) Western blot of HEK293T cells transfected with vectors used for electroporation D) Kaplan-Meier curves of tumor-free survival after electroporation (K+L) E) Exemplary images of mice during the electroporation procedure, upon removal of tumors and bioluminescence imaging. Scale bars equal 5mm. F) H&E histographs exemplifying typical histological appearance of tumors driven by oncogenic *KRAS* and *Trp53* inactivation. Scale bars equal 100µm. G) Agarose gel electrophoresis from tumor gDNA for transposed *RAS* H) TIDE analysis knockout efficiency in tumor compared to wildtype (tail) control tissue.

This optimized protocol resulted in highly efficient tumorigenesis in 2-3 weeks with 100% penetrance and 100% bilateral tumor onset for the *KRAS*/*Trp53* model for both P30 and P0 EPO **(Fig. 1D, S1B-C)**. Resulting tumors displayed sarcoma-like morphology with mostly spindle or mixed spindle/epithelioid features **(Fig. 1E-F)**. Uniformly strong expression of the driving *RAS* oncogene was accompanied by high levels of proliferation marker Ki-67 across all tumors (50-70%). About 70% of tumors showed focal positivity for myogenin indicating some myogenic differentiation **(Fig. 1F)**. As expected, the tumors were positive for the *KRAS* transgene and displayed *Trp53* insertion-deletion mutations (indels) **(Fig. 1G-H)**. Surprisingly, the initially present IVIS signal was often lost during tumor progression **(Fig. S1D)**, despite successful genomic integration of the reporter gene transposons **(Fig. S1E)**. This was particularly evident when reporter genes were expressed from the same vector as the oncogene (KGL) and was accompanied by significantly reduced tumorigenesis **(Fig. S1F)**. This phenomenon suggests immunoediting against reporter genes. Therefore we switched from the outbred CD1 strain, initially chosen due to large litter sizes and good foster qualities, to the inbred C57BL/6J strain which is less susceptible towards immunogenicity to Luciferase and GFP (Han et al., 2008; Skelton et al., 2001). Indeed, no signs of reporter gene silencing were observed in C57BL/6J mice, whereas tumorigenesis efficiency further improved to a mean latency of 14 days (P30) and 21 days (P0) with 100% penetrance and bilateral tumor onset **(Fig. S1G)**. Since SB13 transposase has previously been shown to be limited in transposable cargo size, we further adapted the system to PiggyBac transposase, which showed improved tumorigenesis efficiency for the larger KGL vector in P0 mice **(Fig. S1H-I)**. In summary, we generated an optimized protocol and set of tools to efficiently deliver transgenes and edit tumor suppressors in murine skeletal muscle tissue.

### A versatile genetic toolbox successfully generates several types of fusion-driven sarcoma

Given the high efficiency of the established sarcoma EPO-GEMM system, we next aimed to investigate a broader spectrum of human sarcoma drivers toward their tumorigenic potential **(Fig. 2A)**. For this purpose, we generated a versatile toolbox of transposon and CRISPR vectors with a focus on sarcoma-typical fusion oncogenes, allowing rational combination based on human sarcoma profiling data **(Fig. 2B, S2A-B)**. Reporter genes were omitted to avoid any immunogenicity and maximize tumorigenesis efficiency. Since many fusion-driven sarcomas exhibit few or no co-occurring alterations, oncogenes were tested alone (together with an empty sgRNA vector) and in combination with sgRNAs targeting tumor suppressors, such as *Trp53* to overcome oncogene-induced stress and facilitate cell cycle progression.

**Figure 2.**
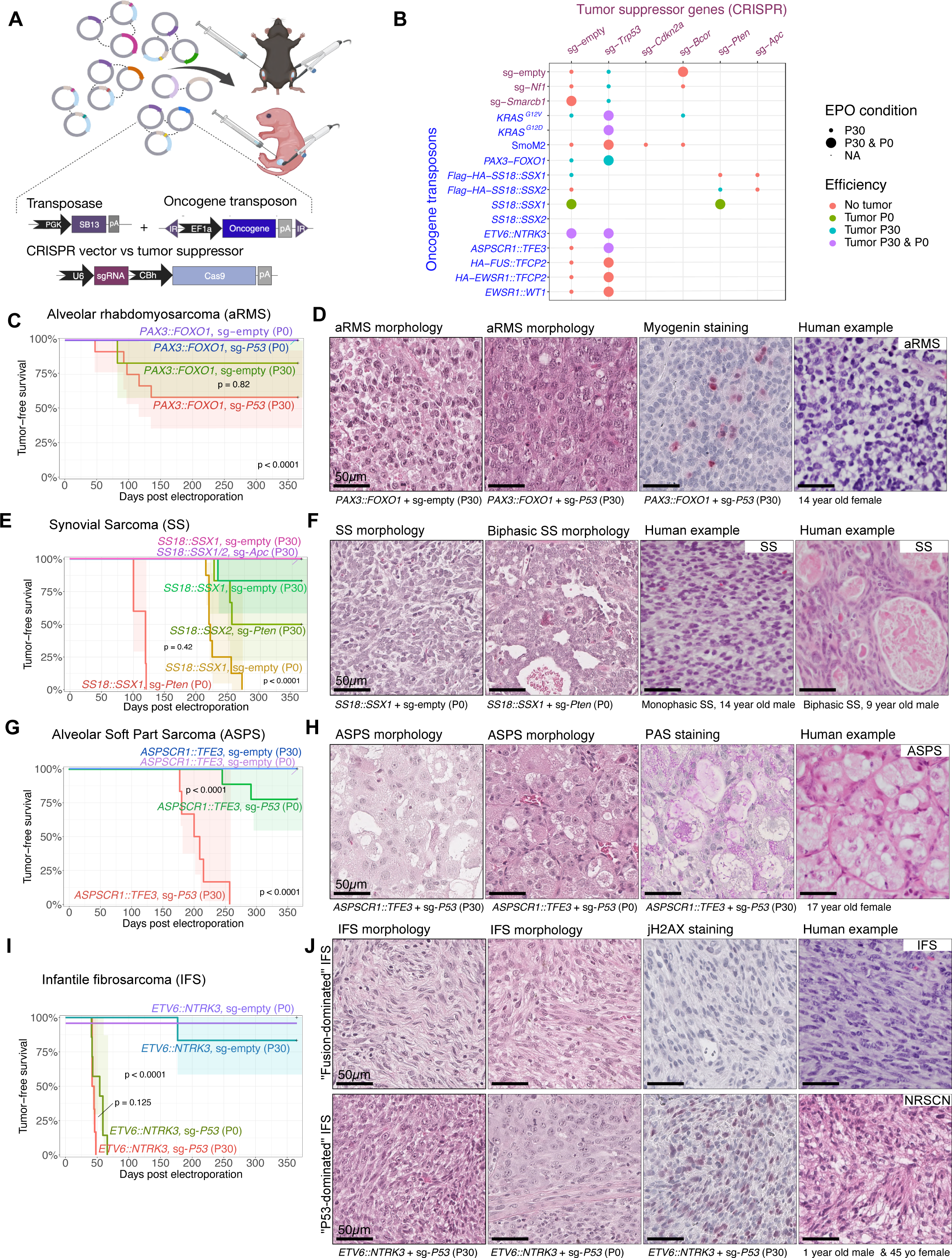
Modeling diverse subtypes of fusion-driven sarcoma A) Scheme of versatile vector combination for *in situ* sarcoma induction. Reporter genes were not utilized here. B) Matrix of vector combinations chosen based on human sarcoma profiling studies. C) Kaplan-Meier curves of murine aRMS induction and representative histographs compared to human aRMS (D). E) Kaplan-Meier curves of murine SS induction and representative histographs compared to human SS (F). G) Kaplan-Meier curves of murine ASPS induction and representative histographs compared to human ASPS (H). I) Kaplan-Meier curves of murine IFS induction and representative histographs compared to human IFS and the emerging WHO entity of NTRK-rearranged spindle cell neoplasm (NRSCN) (J). Murin tumors clustered into two distinct groups, named Fusion-dominated IFS and P53-dominated IFS. If not specified otherwise, histographs were stained by H&E. P-values were determined by log-rank test and corrected for multiple testing with the Bonferroni-Holm method where necessary. Scale bars = 50 µm.

A bona fide example of a clinically aggressive fusion-driven STS is fusion-positive RMS, also known as alveolar RMS (aRMS), which is mainly driven by the *PAX3/7::FOXO1* fusion gene. In our model system, *PAX3::FOXO1*-induced tumorigenesis was more efficient with concurrent inactivation of *Trp53* compared to *PAX3::FOXO1* alone which is in line with previous studies showing that inactivation of *Trp53* or *Ink4a/Arf* critically accelerates aRMS development **(Fig. 2C, S2C-D)** (Keller et al., 2004). *PAX3::FOXO1*/sg*Trp53* tumors occurred unilaterally, with an average tumor-free survival time of 100 days and 40% penetrance rate. Only one of six mice developed a tumor without simultaneous *Trp53* inactivation. Notably, tumors exclusively emerged in P30 mice, but not in P0 animals, suggesting a larger reservoir of cells permissive to PAX3::FOXO1-driven oncogenic transformation in this age group. This corroborates earlier observations in conventional GEMMs that maturing rhabdomyoblasts are more susceptible to PAX3::FOXO1-driven transformation than embryonic or postnatal muscle stem cells (Keller and Capecchi, 2005). Histologically, tumors were highly cellular and mitotically active, consisting of sheets of primitive small blue round cells with hyperchromatic nuclei and small nucleoli, akin to histological features observed in human aRMS (Antonescu et al., 2020) **(Fig. 2D)** and conventional mouse models of this disease (Keller et al., 2004). Myogenin was only partly positive and less pronounced compared to *KRAS/sgTrp53* tumors.

The most common NRSTS in adolescents and young adults, also occurring in children is Synovial Sarcoma (SS) (Sultan et al., 2009). The driving oncofusion *SS18::SSX1* or the less commonly occurring *SS18::SSX2* were electroporated in combination with sgRNAs targeting *Pten* and *Apc*, both of which show loss of function mutations in patient specimens. Notably, SS formation was significantly more efficient in P0 animals where even the fusion alone (+ sg-empty) led to tumorigenesis with 100% penetrance, albeit with mean latencies of about 230 days and 37% bilateral tumor fraction. Inactivation of *Pten* accelerated tumorigenesis to a mean latency of 112 days, 100% penetrance and 100% bilateral tumor fraction **(Fig. 2E, S2C-D)**. Histologically, tumors were highly reminiscent of SS **(Fig. 2F)**. Tumors generally displayed monophasic differentiation, characterized by cellular sheets and at times fascicular growth of uniform spindle cells with scanty cytoplasm and a delicate “chicken-wire” vasculature. A subset of tumors showed a pronounced collagenous stroma with conspicuous bundles of “wiry” collagen. Notably, 53% of tumors demonstrated areas with epithelial, gland-like structures alongside the spindle cell population, which is the characteristic histological feature of biphasic SS in humans (Antonescu et al., 2020). Anti-HA staining confirmed nuclear expression of *SS18::SSX* in tumor cells **(Fig. S2E)**. Overall SS EPO- GEMMs were highly reminiscent the human disease and conventional mouse models (Barrott et al., 2016), further highlighted by nuclear positivity for SS marker TLE-1 (El Beaino et al., 2020) **(Fig. S2F)**.

Another successfully established fusion-driven EPO-GEMM model was Alveolar Soft Part Sarcoma (ASPS), characterized by the *ASPSCR1::TFE3* oncofusion. Similar to aRMS, tumor induction was dependent on *Trp53* inactivation and was more efficient in mice electroporated at P30 rather than at P0 (100% versus 22% penetrance, 33% versus 0% bilateral tumor fraction, 200 versus 269 days of mean latency) (**Fig. 2G, S2C-D**). Human ASPS is characterized by its pathognomonic nest-like, alveolar growth pattern with large, polygonal cells with vesicular nuclei and abundant eosinophilic cytoplasm, typically staining positive in Periodic Acid Staining (PAS). This pathognomonic phenotype, reminiscent of pulmonary alveoli was remarkably well reflected in our model **(Fig. 2H)**, making it indistinguishable from the only hitherto published conventional mouse model (Goodwin et al., 2014) and the human disease (Antonescu et al., 2020).

Lastly, we also applied our new EPO-GEMM system to test gene fusions for which no conventional mouse models have yet been described. A relevant example are *NTRK* translocations, which are considered the characteristic alteration of Infantile Fibrosarcoma (IFS), but also occur in a broad spectrum of malignancies with diverse tissue origin (e.g. soft tissue, brain, thyroid, breast or uterus). NTRK inhibitors have recently been approved as entity-agnostic therapeutics, yet suitable models to optimize treatment to prevent or circumvent resistance mechanisms are largely lacking. To bridge this gap, we delivered the *ETV6::NTRK3* fusion gene alone or in combination with a sgRNA targeting *Trp53*, which led to tumors with 100% penetrance in both P30 and P0 animals, with a moderately higher efficiency in P30 mice (45 days versus 52 of mean latency, 100% versus 0% bilateral tumor fraction). Except for one mouse in the P30 group, tumorigenesis was dependent on *Trp53* inactivation **(Fig. 2I, S2C-D)**. Unsupervised clustering based on gene expression and DNA methylation classified NTRK tumors into two groups **(Fig. 5A-B)**, correlating with distinct histological features **(Fig. 2J)**. 65% (9/14) of tumors showed a fascicular growth pattern consisting of uniform spindle cells with scant cytoplasm and only mild to moderate cytologic atypia. In contrast, 35% (5/14) of tumors showed areas of diffuse growth of spindle to epithelioid cells with a more pronounced eosinophilic cytoplasm and in part (3/5 tumors) severe cytologic atypia, reminiscent of *KRAS/Trp53* tumors. For simplicity, the two groups will be referred to as Fusion-dominated IFS (FD-IFS) and P53-dominated IFS (P53-IFS) hereafter.

Altogether, these results show that the EPO-GEMM approach can be applied to model several gene fusion-driven sarcomas. In each case, the tumors closely recapitulate their human disease counterparts as well as previous conventional GEMMs at the histological level. From all gene fusions tested, only *FUS::TFCP2*, *EWSR1::TFCP2*, and *EWSR1::WT1* did not induce tumors using the current protocol and developmental window. Given the remarkable flexibility of the EPO-GEMM approach, it can be applied to any gene fusion, alone or in combination with relevant sgRNAs, allowing to further optimize tumorigenesis for different subtypes. The successful generation of a mouse model driven by *ETV6::NTRK3* illustrates the potential of this method for many other alterations that have not yet been modeled *in vivo*.

### *Bcor* inactivation cooperates with oncogenic *RAS* to drive fusion-negative RMS

The flexible somatic gene delivery using the EPO-GEMM system allows to probe the cooperativity of different sarcoma-related genetic alterations observed in patient tumors with a short turnaround time. Molecular profiling studies have revealed inactivating, truncating or point mutations of the BCL6 corepressor (*BCOR)* gene, encoding a component of the Polycomb repressive complex 1.1 (PRC1.1), in about 15-20% of eRMS and about 5% of aRMS (Astolfi et al., 2019; Shern et al., 2021). Consistent with a role of BCOR in the context of PRC1.1, *BCOR* mutations are concentrated at the C-terminus containing the PUFD (PCGF Ub-like fold discriminator) domain, which is responsible for interaction with the PRC1.1- defining subunit PCGF1 **(Fig. S3A)** (Shern et al., 2021). Experimental studies have demonstrated a tumor suppressor function of *Bcor* in subsets of leukemia and medulloblastoma (Kutscher et al., 2020; Schaefer et al., 2022), but its role in RMS tumorigenesis remains unclear. We made use of our system to test the hypothesis that *Bcor* inactivation cooperates with oncogenic *RAS* in fusion-negative RMS. A sgRNA directed at exon 3 of *Bcor* resulted in increased tumorigenic efficiency (5/8 mice, 62.5% penetrance) compared to *KRAS^G12V^*alone (1/8 mice, 12.5% penetrance). As expected, the effect was not as strong as with *Trp53* inactivation (100% penetrance) **(Fig. 3A)**. Combination with *PAX3::FOXO1* or inactivation of *Bcor* alone did not lead to tumors (0/8 each). While the histology of *KRAS/sgTrp53* tumors was consistent with patient specimens of *TP53*-mutated eRMS, exhibiting focal or diffuse anaplasia (Pondrom et al., 2020), *KRAS/sgBcor* tumors were predominantly monomorphic with few signs of pleomorphism and showed focal rhabdomyoblastic differentiation, consistent with *TP53* wildtype eRMS morphology **(Fig. 3A)**. This was reflected in a lower rate of genomic instability and DNA double strand breaks in *KRAS/sgBcor* tumors when compared to *KRAS/sgTrp53* **(Fig. 3E-F).** Interestingly, in contrast to *KRAS/sgTrp53* tumors, *KRAS/sgBcor* tumors were negative for mesenchymal markers desmin and myogenin **(Fig. S3B)** and showed a decreased average DNA methylation, possibly indicative of a different cell of origin, differentiation state or as a result of epigenetic reprogramming during the transformation process **(Fig. S3C-D)**. Taken together, these results provide experimental confirmation of *Bcor’s* role as a tumor suppressor gene in eRMS.

**Figure 3.**
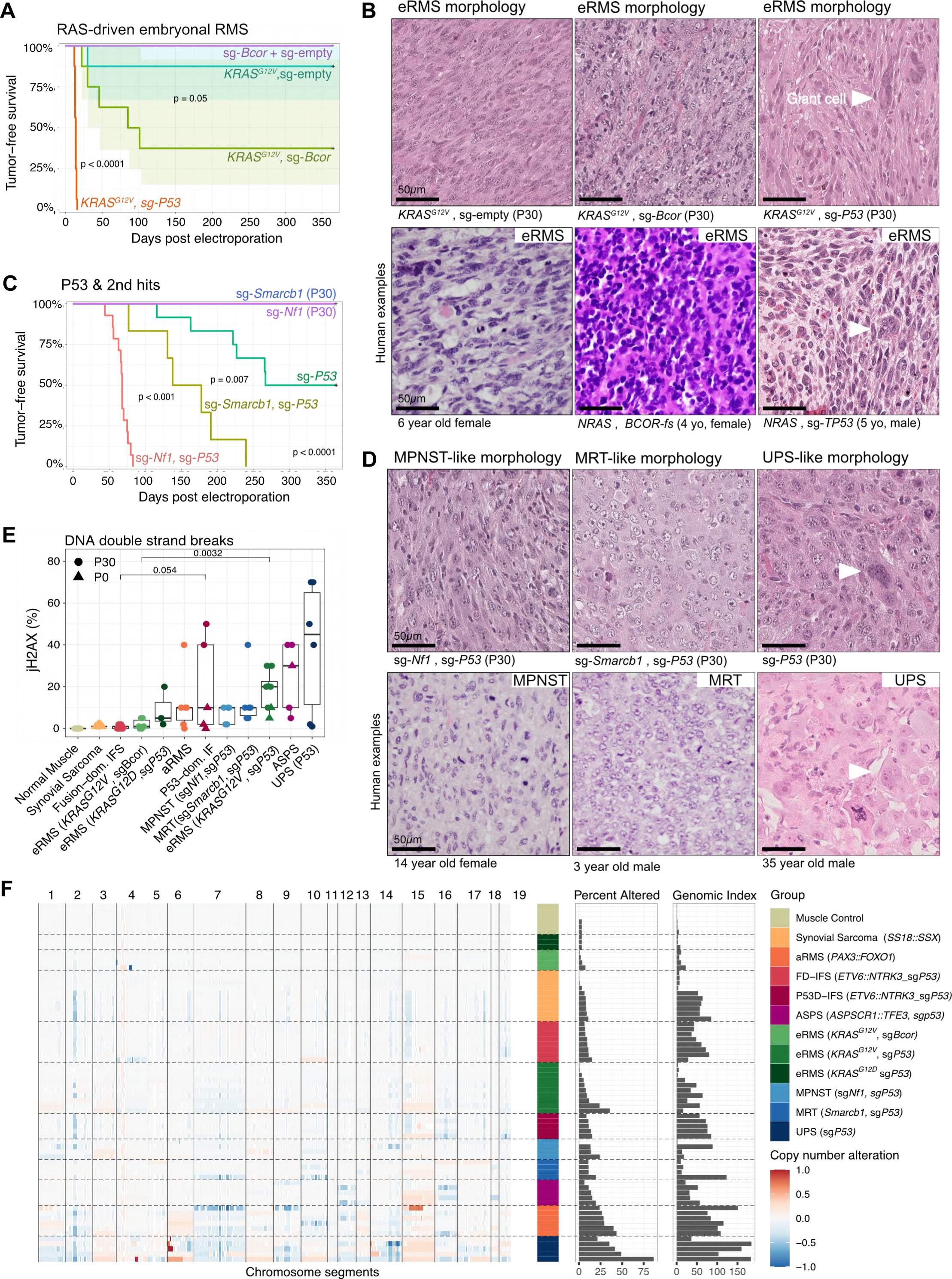
Modeling sarcomas driven by oncogenic RAS and gene inactivation. A) Kaplan-Meier curves of *RAS*-driven mouse sarcomas and representative H&E histographs compared to human eRMS (B). C) Kaplan-Meier curves of mouse sarcomas driven by *Trp53*- inactivation and secondary mutations and representative histographs compared to human MPNST, MRT and UPS (D). E) Integrated score for jH2AX IHC stainings, depicted as boxplots ordered by median from low to high. P-values of boxplots were determined by unpaired two-sided Wilcoxon tests and corrected for multiple testing by Bonferroni-Holm method. F) CNV profiles derived from DNA methylation data, condensed as Percent altered (≤/≥ .1) and Genomic Index. N ≥ 3 tumors per group. Scale bars = 50 µm.

### *Trp53* inactivation cooperates with second hit mutations to drive a spectrum of pleomorphic sarcomas

CRISPR-mediated inactivation of *Trp53* alone led to unilateral tumor formation in 50% of cases with a mean latency of 210 days **(Fig. 3C)**, essentially mimicking pleomorphic sarcoma formation in germline-mediated mouse models of Li-Fraumeni syndrome (Lang et al., 2004; Olive et al., 2004). While the models described above each have clearly delineated oncogenic drivers, *Trp53*-mediated genomic instability and diachronous acquisition of second hit mutations over time are the likely cause of oncogenic transformation in this context (Gerstung et al., 2020). This was reflected in pronounced copy number variations (CNVs) **(Fig. 3F)** and the highest rate of DNA double strand breaks across the sarcoma EPO-GEMM cohort (**Fig. 3E)**, qualifying this model system to study the nature of mut*Trp53*-mediated genome evolution in the future (Baslan et al., 2022). Synchronous *Smarcb1* inactivation, pathognomonic for Malignant Rhabdoid Tumors (MRT) and Epithelioid Sarcomas (EpS), accelerated tumor formation to a mean latency of 173 days with 100% penetrance and 17% bilateral tumor fraction, which is significantly more efficient compared to a previously reported Myf5-Cre-mediated Smarcb1-inactivation model exhibiting 40% and latency longer than 12 months (Li et al., 2021). The efficiency of tumorigenesis was even higher upon synchronous *Nf1* loss, conveying indirect activation of RAS/MAPK signaling **(Fig. 3C)**, typically observed in patients suffering from Neurofibromatosis Type 1 (NF-1), who exhibit an increased risk for malignant peripheral nerve sheath tumors (MPNST) or *Nf1*-deleted eRMS (Maude et al., 2018). Combined *Nf1/Trp53* inactivation, yielded tumors with a mean latency of 68 days, 100% penetrance and 86% bilateral tumor fraction, exceeding the efficiency of a previously reported CRISPR-mediated somatic mouse model of combined *Nf1/Trp53* inactivation which exhibited a median latency of approximately 100 days (Huang et al., 2017).

Histologically, tumors from all three groups were reminiscent of their human counterparts. sg*Nf1/*sg*Trp53* tumors were compatible with MPNST or *Nf1*-inactivated eRMS with a fascicular growth pattern of spindle or spindle/epithelioid cells **(Fig. 3D)**. The S100 protein neuronal marker was only focally positive in 3/5 tumors, while the myogenic marker myogenin showed stronger focal positivity in 4/5 tumors, suggesting a predominantly myogenic differentiation **(Fig. 4A)**. sg*Smarcb1/*sg*Trp53* tumors consisted of sheets of large epithelioid tumor cells with vesicular and eccentric nuclei amidst an abundant eosinophilic cytoplasm, reminiscent of the characteristic rhabdoid cell morphology of human MRT or EpS **(Fig. 3D)**. While highly positive for mesenchymal marker desmin, they were negative for myogenin and heterogeneous in the expression of other differentiation markers such as ASMA, cytokeratin, and S100 protein **(Fig. 4A)**. Murine UPS (sg*Trp53*-only tumors) were strongly positive for desmin and showed heterogeneous expression of other differentiation markers **(Fig. 4A)**. Anaplasia was noted in all three groups, but was most pronounced in sg*Trp53-*only tumors (sg*Trp53* > sg*Smarcb1/sgTrp53* > sg*Nf1/sgTrp53*) where giant cells and atypical mitoses were also frequently found **(Fig. 3D)**. In conclusion, the sarcoma EPO- GEMM system provides an excellent platform to study *Trp53*-mediated genome evolution and pleomorphic sarcomas driven by tumor suppressor gene inactivation.

**Figure 4.**
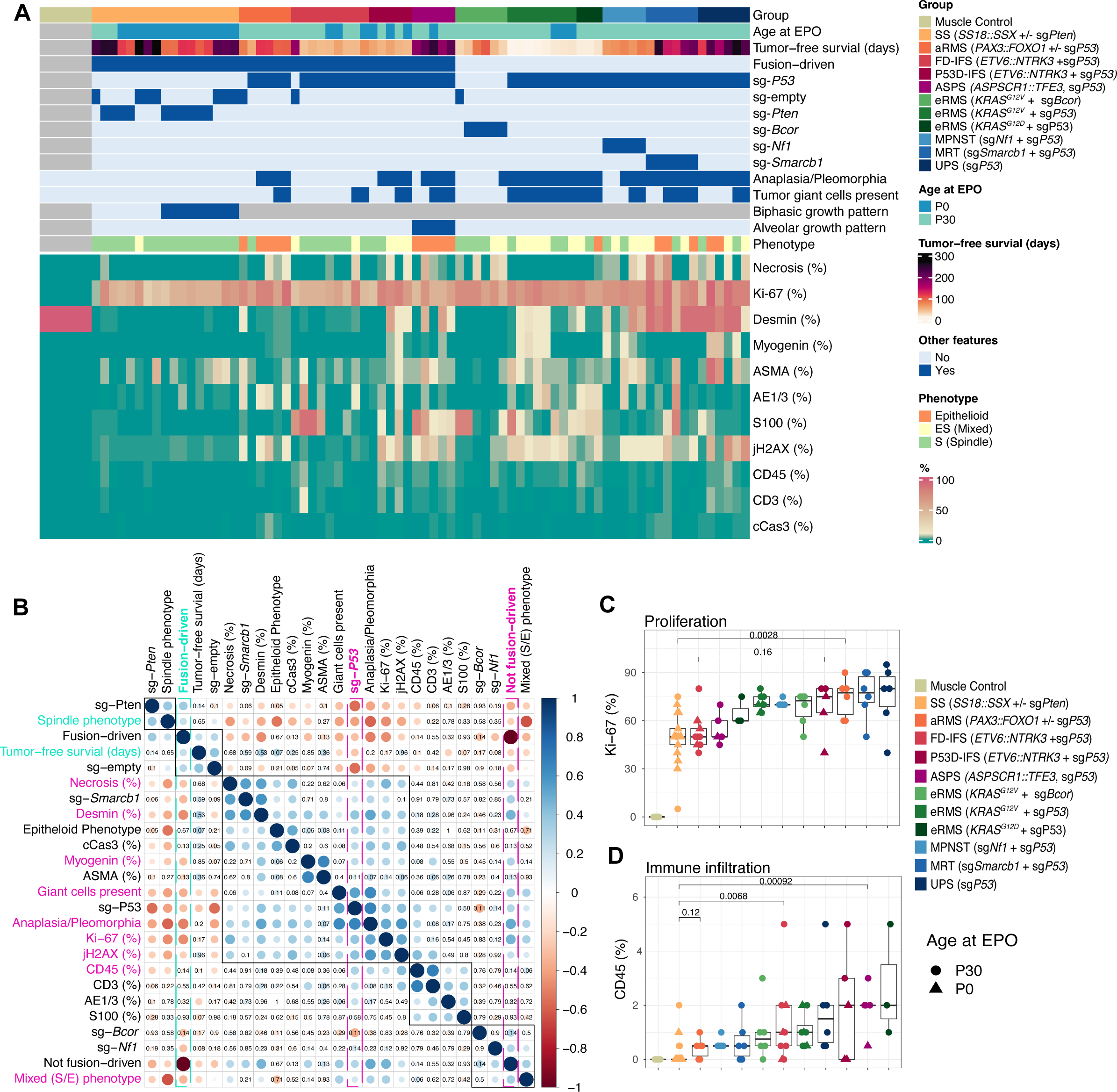
Fusion gene and TP53 mutation status determine sarcoma biology and microenvironment A) Heatmap view of mouse sarcomas, quantified for six morphological and based on H&E staining and integrated expression scores for 10 antigens determined by immunohistochemistry. Asymmetric color scale for combined visualization of low (CD45, CD3, cCas3) and high-scoring antigens. Quantification by blinded expert pathology review. B) Pearson correlation matrix based on data from panel A. Insignificant p-values (≥ 0.05) were plotted. Blue indicates positive, red negative correlation, point size reflects effect size. C-D) IHC scores of Ki-67 and CD45 from panel A visualized as boxplots ordered by median from low to high. P-values of boxplots were determined by unpaired two-sided Wilcoxon tests and corrected for multiple testing by Bonferroni-Holm method. N ≥ 3 tumors per group.

### Fusion gene and *Trp53* status determine sarcoma biology and microenvironment

Given the unique opportunity to systematically compare a large number of different sarcoma types established under identical genetic and experimental background, we performed an in- depth assessment of their histological phenotypes. A blinded expert pathology review of H&E stains and ten IHC markers was performed across the mouse sarcoma cohort, systematically assessing signs of anaplasia, growth pattern and cellular phenotype as well as immunoreactivity **(Fig. 4A-B)**. While necrosis was generally low except for MRT, all models were highly proliferative as quantified by Ki-67 staining. aRMS, UPS and MRT showed particularly high proliferation rates in line with their human counterparts **(Fig. 4C)** (Antonescu et al., 2020).

Correlation analysis of quantified features **(Fig. 4B)** confirmed that fusion gene and *Trp53* mutation status are major determinants of sarcoma biology. Compared to non-fusion-driven models, fusion-driven sarcomas typically consisted of rather uniform, spindle cells. In these tumors, occurrences of anaplasia and tumor giant cells were rarely observed. They were also characterized by lower proliferation (Ki-67) and DNA double strand break rates (phosphorylated H2AX/jH2AX) and reduced immune infiltrates as quantified by CD45 staining. Exceptions were the predominantly small round cell, but likewise uniform growth pattern in aRMS and the higher immune infiltrate in ASPS, also reported for the human disease (Brohl et al., 2021). In contrast, and as observed in human tumors (Demicco et al., 2017), non-fusion driven models frequently consisted of irregular and spindle to epithelioid tumor cells, frequently exhibiting anaplasia, as well as higher rates of DNA double strand breaks, proliferation and immune infiltration (**Fig. 4B-D**). Importantly, the positive correlation between genomic stability and immune infiltration found in human sarcoma cohorts, was preserved across sarcoma EPO-GEMMs (Tazzari et al., 2021). Although the general degree of immune infiltration was rather low, as to be expected for sarcomas, leukocyte aggregations up to tertiary lymphocytic structures were occasionally observed **(Fig. S4A-B)**.

To further validate the immunophenotype observed by IHC, we employed CibersortX as an algorithm for immune cell deconvolution from bulk RNA sequencing of tumors **(Fig. S5A- B)**. There was a modest but statistically significant correlation for the total leukocyte fraction across IHC and CibersortX **(Fig. S5C)**, which is likely due to the sampling bias for bulk RNA seq and the overall low immune infiltration observed across sarcoma subtypes. Whereas both SS and aRMS were noticeably immune-cold across both methods and across further immune cell subpopulations, RNA-sequencing-based deconvolution failed to detect increased immune infiltration in ASPS **(Fig. S5D-G)**.

Of note, EPO-GEMM primary tumors and derived cell lines, recapitulated the expression patterns of various immunotherapy targets antigens **(Fig. S6A-F)**, currently explored for human sarcoma treatment (Baldauf et al., 2018). The immune checkpoint surface marker PD- L1 was not upregulated compared to muscle controls in most tumors, indicating absence of immunoediting **(Fig. S6A)**. However, several pan-cancer immunotherapy target antigens including B7-H3/CD276, GD2 and Erbb2/HER2/neu were markedly upregulated in sarcoma GEMMs **(Fig. S6B-D)**. Particularly B7-H3 was strongly upregulated across the entire spectrum of sarcoma EPO-GEMMs **(Fig. S6B and F)**, further supporting ongoing clinical approaches of anti-B7-H3 immunotherapy in solid tumors. Additionally, some target antigens were upregulated in subtype-specific fashion, including MCAM (Melanoma cell adhesion molecule) in FD-IFS or FGFR4 in aRMS, both of which are currently being explored as immunotherapy approaches in Malignant Melanoma (Rapanotti et al., 2021) and aRMS (Sullivan et al., 2022) respectively. Taken together, unsupervised molecular and histological analyses of sarcoma EPO-GEMMs confirmed fusion gene status and *Trp53* mutation status as the major determinants of sarcoma biology.

### Mouse sarcomas exhibit distinct genotype-dependent molecular signatures

Unsupervised tumor classification based on DNA methylation profiling has significantly expanded the means to accurately classify human brain tumors (Capper et al., 2018) and sarcomas (Koelsche et al., 2021). We adopted this approach to our sarcoma EPO-GEMM cohort by using the recently released 285k mouse methylation array (Zhou et al., 2022). Clustering of EPO-GEMM tumors based on all or the top 10,000 differentially methylated probes led to genotype-dependent clustering, which was particularly evident for the fusion-driven tumor types aRMS, SS, ASPS and FD-IFS **(Fig. 5A, S3D)**. Consistent with overlapping microscopic features, non-fusion driven tumors and P53D-IFS exhibited a more diffuse distribution compared to fusion-driven models. Tumor specimens from a conventional mouse model of SS where *SS18::SSX2* is expressed from the *Rosa26* locus upon Cre recombination (Barrott et al., 2016), clustered together with SS EPO-GEMMs, indicating conservation of SS18::SSX-mediated biology between these two modeling approaches **(Fig. 5A, S3D)**. One notable phenomenon observed was a broad hypomethylation phenotype in the Synovial Sarcoma group **(Fig. 3C-D)** as previously demonstrated by (Demicco et al., 2017) for human SS. This is consistent with the hypothesis that the SS18::SSX fusion mediates epigenetic rewiring and oncogenic transformation through binding to unmethylated CpG islands (Banito et al., 2018). Tumors from a previously published conventional GEMM of ASPS, expressing *ASPSCR1::TFE3* from the *Rosa26* locus, exhibited distinct DNA methylation profiles **(Fig. 5A, S3D)**, possibly reflecting a different cell of origin, given their strictly heterotopic onset in the cranial vault (Goodwin et al., 2014).

**Figure 5.**
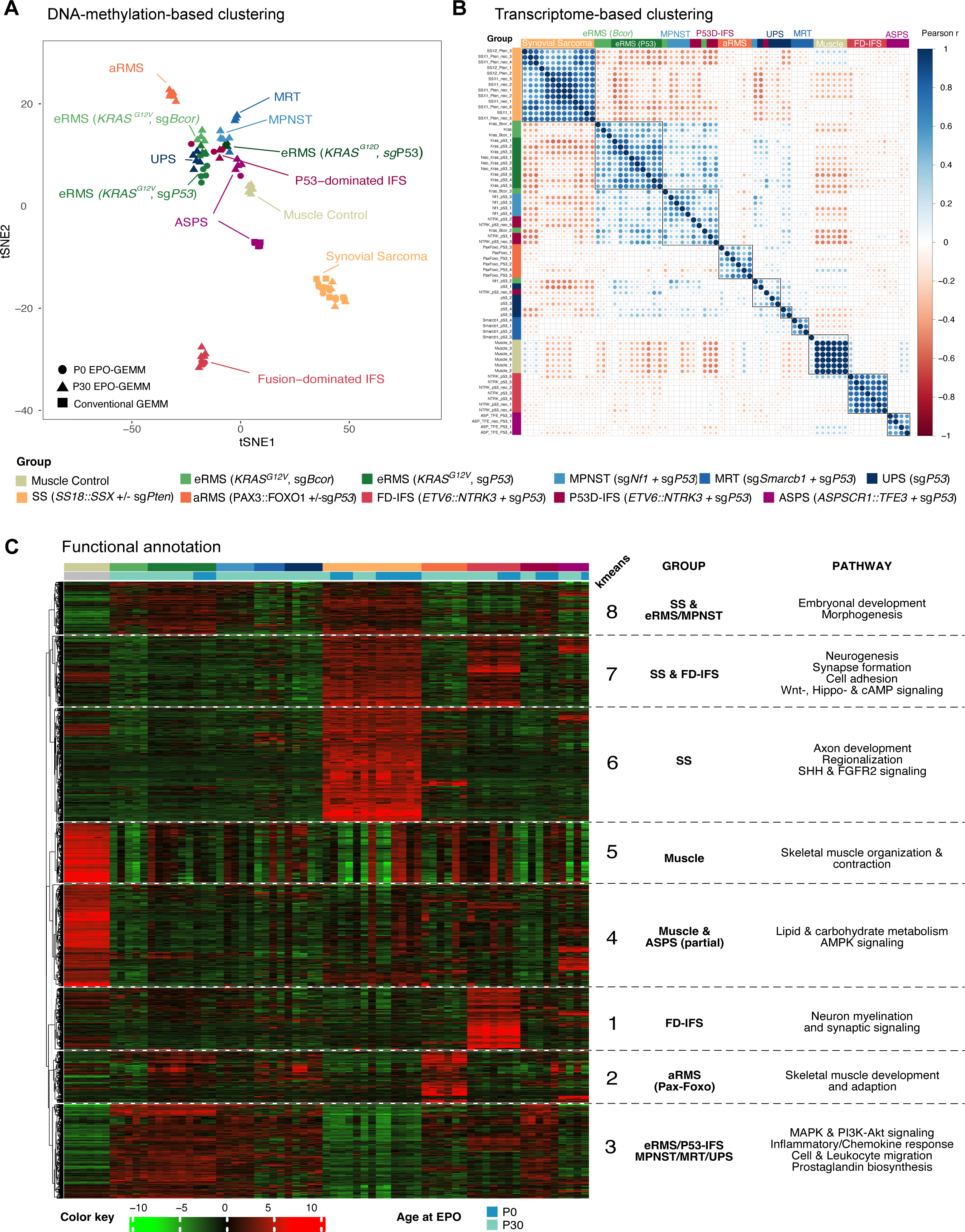
Mouse sarcomas exhibit distinct genotype-dependent molecular signatures **A)** tSNE clustering based on the top 10,000 differentially methylated CpG sites. **B)** Correlation clustering based on the top 2,000 differentially expressed genes. **C)** k-means clustering based on the top 2,000 differentially expressed genes, annotated with summarized GSEA data (Gene set enrichment analyses) from ten gene set collections, i.e. GO Biological Process, GO Molecular function, GO cellular component, KEGG, Reactome, Wiki pathways, Jenssen Compartments, Jenssen Tissues, GeneSetDB and TF Regulatory Networks.

Transcriptome-based clustering corroborated these results with fusion-driven mouse sarcomas forming distinct clusters each corresponding to genotype-specific expression patterns **(Fig. 5B)**. *KRAS/sgTrp53/sgBcor* and sg*Nf1/sgTrp53* models showed fairly similar transcriptomes, consistent with the commonly underlying upregulation of RAS/MAPK signaling **(Fig. 5C)**. While FD-IFS formed a very distinct cluster, PD53-IFS clustered in a rather scattered fashion adjacent to eRMS and UPS tumors. To further explore the biology underlying induced murine sarcomas, RNA sequencing data was further subjected to k-means clustering and Gene Set Enrichment Analysis (GSEA), using ten different gene set collections from the Molecular Signature Database (MSigDB) **(Fig. 5C, Table S2)**. As expected, muscle control tissue (k-means clusters 4 and 5) was clearly distinct from all tumor types and identified genes related to skeletal muscle organization and contraction. Upregulation of embryonal developmental pathways (k-means group 8) was shared between various tumor types while muscle development and adaptation were specifically upregulated in aRMS, consistent with mechanistic studies identifying *PAX3/7::FOXO1*-induced activation of myogenic super enhancers (Gryder et al., 2017). Consistent with the underlying *NTRK* fusion, signaling pathways of neuron myelination and synapse signaling were specifically upregulated in FD-IFS (k-means group 1), whereas enrichment of neuro- and synaptogenesis, Wnt, Hippo, and cAMP signaling were shared with SS (k-means group 7). SS displayed a unique expression signature (k-means group 6), including developmental pathways known to be enriched in human tumors, such as axon development, Sonic Hedgehog (SHH) and FGFR2 signaling (Banito et al., 2018). ASPS shared signatures with other entities which included upregulation of genes involved in lipid and carbohydrate metabolism, consistent with findings in a previous conventional mouse model of ASPS (Goodwin et al., 2014).

These results vividly demonstrate how the diversity of underlying genetic perturbations drives transcriptomically diverse tumors with a clear genotype-phenotype association. Particularly, sarcoma-driving oncofusions elicit profoundly specific tumor transcriptomes that reflect their respective underlying oncogenic mechanisms.

### Murine sarcomas faithfully recapitulate the human sarcoma spectrum

To systematically assess whether sarcoma EPO-GEMMs recapitulate their human counterparts, we used an unbiased cross-species bioinformatic approach with correction for species and dataset **(Fig. 6A)**. To represent all relevant human sarcoma subtypes, we integrated and harmonized RNA sequencing data from the ‘The Cancer Genome Atlas’ (TCGA) Sarcoma study (Demicco et al., 2017), St. Jude Cloud (McLeod et al., 2021), and the INFORM registry (van Tilburg et al., 2021), yielding a total of 397 human sarcoma samples that were compared to 63 mouse sarcoma samples. After restricting genes to cross-species-conserved orthologues, differential gene expression analysis between each entity versus all, and each entity versus control tissue was performed for both human and murine datasets. The resulting list of 2636 genes that were present in both mouse and human genomes (**Table S3**) was batch-corrected for species effects and used for hierarchical clustering **(Fig. 6B).** Mouse sarcomas, in particularly bona fide fusion-driven sarcomas, clustered together with their human counterparts.

**Figure 6.**
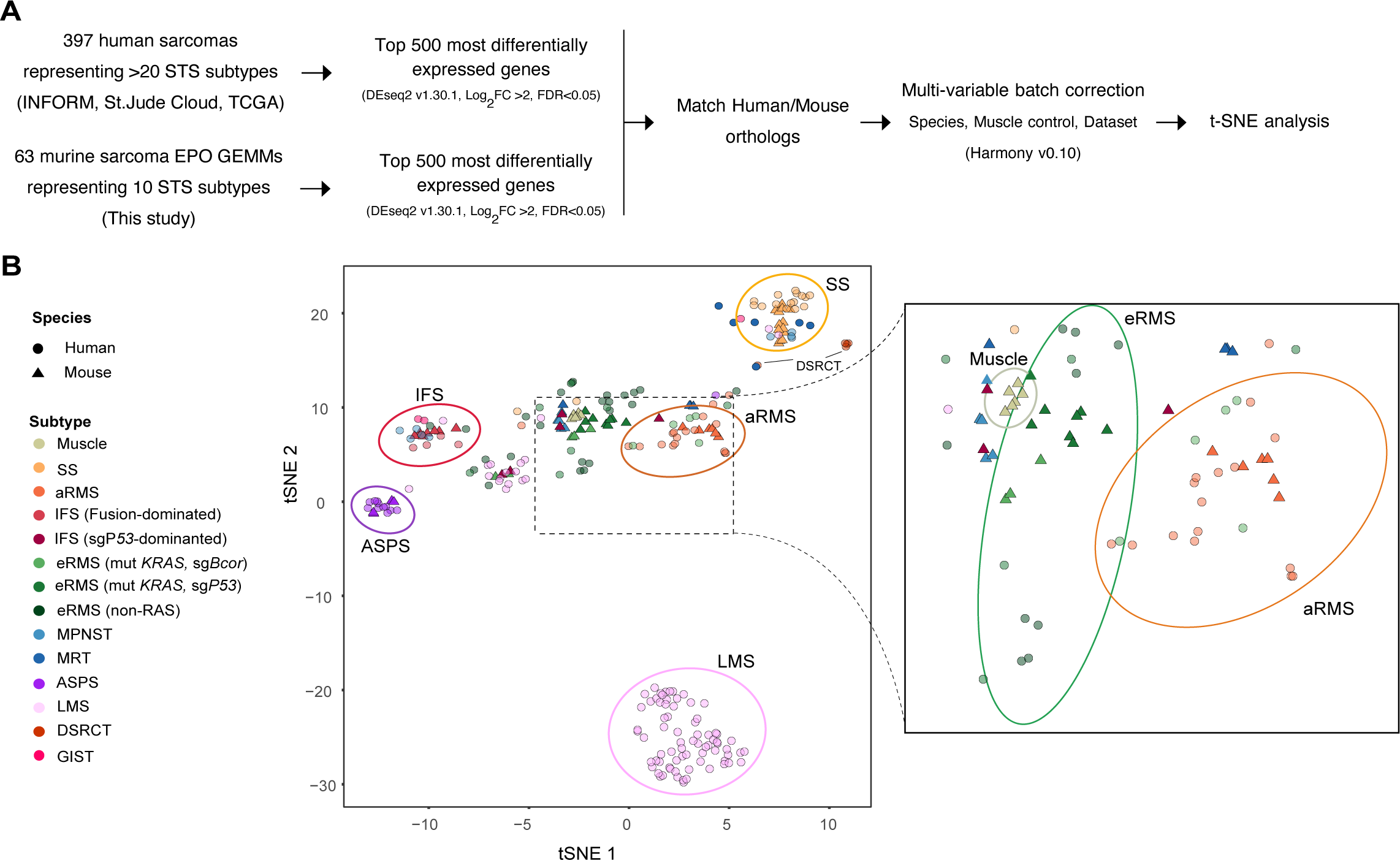
Murine sarcomas faithfully resemble the human sarcoma spectrum A) Analysis scheme of cross-species sarcoma analysis based on RNA sequencing. B) tSNE clustering based on cross-species transcriptome analysis of mouse (n=63) and human (n=397) sarcoma specimens.

Overall, cross-species comparison of transcriptome, methylome, and histology data revealed a coherent picture, indicating conservation of human sarcoma biology across the sarcoma EPO-GEMM cohort, especially for bona-fide fusion-driven sarcomas SS, FD-IFS, aRMS and ASPS.

### Syngeneic allograft models preserve GEMMs for long-term application

In terms of therapeutic predictive value, both patient derived xenografts (PDX) models and GEMMs (Stewart et al., 2017) are regarded as superior when compared to conventional human cancer cell lines which are still used for about 80% of preclinical therapy trials (Gengenbacher et al., 2017; Stewart et al., 2017) Most GEMMs have predominantly been used to study the pathobiology of tumors, leaving their translational potential vastly unexplored. The typically mixed genetic backgrounds of conventional GEMMs are hampering immunocompetent allografting and cross-institutional model sharing. Hence, the broad panel of genetically heterogeneous sarcoma GEMMs established here on genetically identical C57BL/6J background provided an excellent opportunity to explore the appropriate methodology to systematically preserve GEMMs for preclinical treatment trials.

First we tested whether primary tissue allografts would be superior to *in-vitro-*propagated cells (Jung et al., 2018) and tumor spheres superior to conventional 2D cultures (LeSavage et al., 2022) in preserving the biological properties of the original tumor tissue. Four different syngeneic allograft model (SAM) types were compared upon orthotopic unilateral engraftment: dissociated primary cells (C), 50-200µm tumor fragments (F), cells cultured *in vitro* under 2D conditions (2D) and tumor spheres cultured *in vitro* under 3D serum-free conditions (3D) **(Fig. 7A)**. All four methods substantially reduced mean TFS from about 78 to 15 days across a variety of genotypes **(Fig. 7B, S7A)** with engraftment success rates of about 90%, which is significantly higher than in PDX models (Stewart et al., 2017). Blinded expert pathology review of H&E and IHC sections revealed accurate histomorphology preservation **(Fig. S7B)**. While necrosis and proliferation rates were slightly increased, no signs of immune rejection could be observed in any of the four SAM methods tested **(Fig. S7C-E)**. Importantly, all SAMs and cell lines clustered together with their respective GEMM upon DNA methylation analysis and showed remarkably high conservation of the underlying methylome features **(Fig. 7C-D)**. Broad hypomethylation in SS and aRMS for example was highly preserved across all SAM types as were CNV profiles **(Fig. S7F-G)**. To our surprise, none of the four methods emerged superior in preservation of histotype, genomic stability and DNA methylome as all models recapitulated the original molecular makeup remarkably well. Even 2D-cultured allografts reflected the distinct histomorphology of fusion-driven sarcoma GEMMs **(Fig. 7E)**. In summary, both primary and cell culture-based allograft methods were well suited for the preservation of sarcoma EPO-GEMMs for preclinical testing.

**Figure 7.**
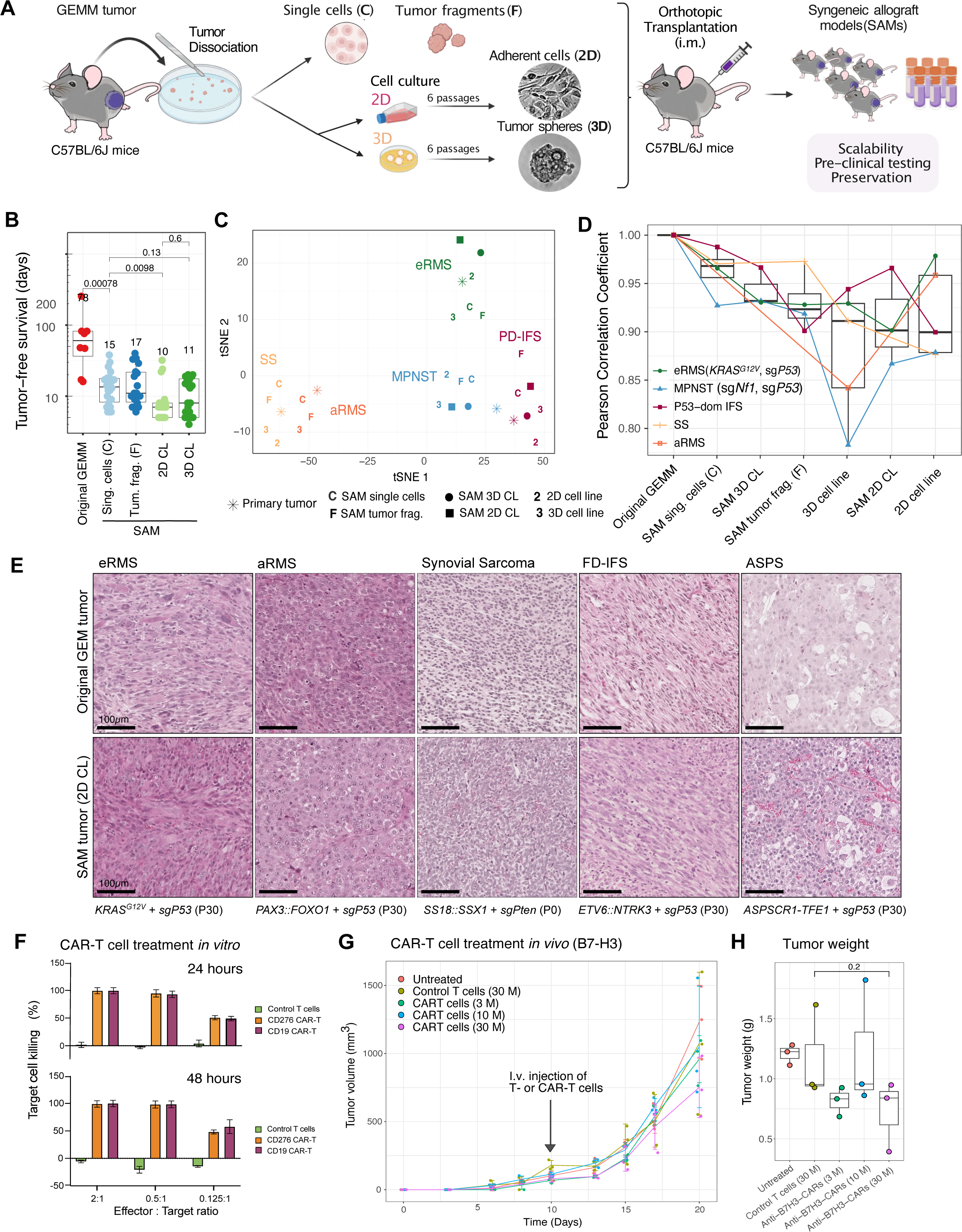
Syngeneic allograft models enable scalability for *in vivo* testing **A)** Schematics of Syngeneic Allograft Modeling (SAM) procedure, systematically comparing four different allograft types. **B)** Tumor-free survival of SAMs compared to corresponding GEMMs. **C)** tSNE clustering based on the top 10,000 differentially methylated CpG sites of SAMs, mouse-tumor-derived cell lines and corresponding GEMMs. **D)** Median-ordered boxplots of Pearson correlation coefficients based on DNA-methylation data from GEMMs and corresponding SAMs. E) Representative H&E histographs of GEMMs and corresponding SAMs after on orthotopic engraftment of 2D-cultured mouse tumor lines. Scale bars equal 100µm. F) *In vitro* co-culture assay of murine aRMS tumor cells and Anti- hCD19 and Anti-hB7-H3-CAR-T cells or untransduced control T cells for 24 and 48 hours. **G)** Tumor volume of orthotopically allografted aRMS tumors, treated i.v. with CAR-T cells (3×10^6^ / 10×10^6^ / 30×10^6^ per mouse) or untransduced control T cells (30×10^6^), compared to untreated control group. I.v. injection of T cells was performed on day 10 after tumor allografting. **F)** Boxplots of weights (g) of extracted tumors 20 days post engraftment. P- values refer to unpaired two-sided t-test. N=3 mice per group.

Finally, we leveraged the flexibility of *in vitro* propagation and syngeneic *in vivo* allografting of sarcoma EPO-GEMMs to apply some of our models to small molecule and immunotherapy applications. NTRK inhibitor therapy with first (Larotrectinib) and second generation (Repotrectinib) agents showed specific and significant activity in FD-IFS and P53D-IFS lines, comparable to response rates in a rare patient-derived cell model acquired from a patient suffering from NTRK-driven Inflammatory Myofibroblastic Tumor (IMT) **(Fig. S8A-B)**.

We also exploited our system to assess immunotherapy approaches by *in vitro* modifying a murine *PAX3::FOXO1*-driven EPO-GEMM sarcoma cell line to express luciferase and human B7-H3 and CD19 as target antigens **(Fig. S8C)**. Treatment with murine CAR-T engineered to detect either human B7-H3 or human CD19 showed highly efficient tumor cell killing *in vitro* **(Fig. 7F, S7D)**. *In vivo* treatment of the same tumor model resulted in only moderate dose-dependent anti-tumor responses **(Fig. 7G-H)**, indicative of immune escape, well known to be hampering cellular immunotherapy approaches for solid tumors in humans. Thus, this translationally relevant model system is excellently suited to optimize cellular immunotherapies for solid tumors in the future, thereby addressing a critical and unmet clinical need in the field of immunotherapy.

## DISCUSSION

The lack of sarcoma models has hindered our understanding of sarcomagenesis and the development and testing of new therapeutic approaches for patients over the last decades. Here we describe an efficient and highly versatile approach to probe any genetic alterations by *in vivo* somatic engineering of mouse muscle tissue. Besides the dramatic reduction in animal numbers by avoiding extensive intercrossing, there are several advantages to EPO- GEMMs when compared to previously derived sarcoma models. Perhaps the most pertinent is that it bypasses the need for conditional transgenic lines and therefore a predetermined assumption of a suitable linage or cellular background for transformation. While Cre- transgenic mouse lines have been elegantly used in the past to show different outcomes when certain gene fusions are expressed in various cellular backgrounds (Abraham et al., 2014; Haldar et al., 2007), EPO-GEMMs or in alternative the delivery of exogenous Cre (e.g. TAT- Cre injection), allow to hit any cell of interest in a given tissue, contingent that the transfection efficiency is high enough. This was a key step in the successful establishment of these models. We substantially optimized muscle electroporation in P30 mice compared to previous studies (Huang et al., 2017; Young and Dean, 2015), but also adapted the procedure to apply it for the first time in neonatal mice, marking the earliest murine soft-tissue electroporation to date. Of note, *in utero* EPO can be applied to brain tissue, but relies on injections into the lumen of cerebral ventricles (Feng et al., 2018), a structural feature not present in other tissues. In assessing the conditions for the highest transfection efficiency of muscle tissue at two postnatal developmental time points, we increased the probability of hitting cells permissive for transformation even if they are rare. This is particularly relevant for gene-fusion driven sarcomas where a precise initial cellular state is thought to be required to allow oncogenic transformation (Davis and Meltzer, 2007). Indeed, our approach could be applied to several gene fusion sarcomas, each exhibiting a defined DNA methylation profile most probably primarily reflective of alternative cells of origin. One could argue that as long as a given cell of origin is present at the time of tissue transfection, the method can be applied to any genetic alteration in the future. Still, there are limitations. Some oncogenes did not give rise to tumors within the observation period of one year post electroporation. These included *TFCP2* fusions (*FUS::TFCP2* and *EWSR1::TFCP2*) and *EWSR1::WT1*, alone or in combination with *Trp53* inactivation. Given that sg*Trp53* alone eventually led to tumors with about 50% penetrance, one would expect similar rates when combining sg*Trp53* with these oncogenes. However, the actually observed rates of sg*Trp53*-only tumors were significantly lower (*TFCP2*-fusions: 2/24 (8%); *EWSR1::WT1* fusion: 1/12 (8%), indicating that most of the transfected cells were probably cleared due to oncogene-induced stress, cell death or immune-related responses. Precisely why these alterations were unsuccessful in generating tumors could be due to a variety of reasons. One of the most important certainly is availability of permissive cells of origin. DSRCT, driven by *EWSR1::WT1,* presents mostly in the abdomen, possibly originating from mesothelial (Tsoukalas et al., 2020) or other non- myogenic cells (Wu et al., 2022). Similarly, TFCP2-fused RMS typically occur intraosseusly or are bone-associated, suggesting a non-myogenic origin (Schöpf et al., 2024). For some entity-specific alterations, embryonic cell states may be required for oncogenic transformation, as previously described for induction of brain tumors by in utero electroporation (IUE) (Feng et al., 2018). This likely explains why *Smarcb1*-inactivation alone was not sufficient for tumorigenesis in our model system, and concomitant *Trp53*- inactivation was required to drive EpS/MRT-like tumors (Han et al., 2016; Li et al., 2021). For some subtypes one cannot rule out the need to co-deliver additional alterations to enable tumorigenesis or the requirement to finetune oncogene expression levels by the use of strength-variable or endogenous promoters. Finally, some oncogenes may never work simply due to functional divergence between human and mouse genomes. Nevertheless, the fact that it was successful for most entities, including the first IFS model and orthotopic ASPS model outside the cranial vault (Goodwin et al., 2014), clearly suggests that this method is applicable to many other sarcoma subtypes.

The procedure also allows multiplexing different genetic alterations and can be applied to mice of any genetic background. An obvious application is to test cooperation between genetic events. In RMS, *BCOR* harbors inactivating mutations in up to 15-20% of cases being the second most frequent mutated gene in this disease after *NRAS* (Shern et al., 2021). Our approach was able to show for the first time that *Bcor* inactivation enables *KRAS^G12V^-*driven tumorigenesis in the muscle. The tumors exhibited reduced expression of myogenic markers, and lower overall DNA methylation when compared to *KRAS^G12V^*/sg*Trp53* tumors, arguing in favor of a *Bcor* inactivated-specific phenotype related to induction of permissiveness in alternative cells of origin and/or to an active epigenetic reprograming that favors transformation. Further studies are needed to mechanistically understand the role of BCOR loss of function in RMS and other pediatric tumors where it is frequently observed (Astolfi et al., 2019).

Overall, this flexible approach allowed to model a very heterogeneous and diverse spectrum of sarcomas. It generated a genetically diverse set of subtypes that are clinically relevant and span from fusion-driven uniform and immune cold sarcomas to more pleiomorphic, genetically unstable tumors, with a tendency towards increased immune infiltrates, dominated by *TP53* mutations. Most importantly, the somatically induced sarcoma models were faithful representatives of previously established conventional GEMMs and the human disease. They reproduced entity-specific histologies with remarkable accuracy and activated expression programs that reflect the biology of the human disease. Another key point of this study was to preserve the variety of new immunocompetent mouse models for use in preclinical treatment studies. Re-engraftment of GEMM tumor material into syngeneic C57BL/6J mice was very efficient without signs of immune rejection. Systematic comparison of four different model preservation methods demonstrated that both primary as well as cell-culture based allografting was suitable for EPO-GEMM preservation and expansion. It should be noted however that we did not perform a thorough analysis of immune infiltration profiles in these syngeneic models and previous studies showed distinct immune landscapes in autochthonous sarcomas and their corresponding transplants (Wisdom et al., 2020). Still, syngeneic models are critical not only for preservation but to allow the scalability required for preclinical studies. This was well demonstrated by their use in recapitulating barriers to efficient CAR-T cell activity *in vivo*.

In summary, our study meets a crucial gap in sarcoma and solid tumor research. It not only presents a suite of models ready for immediate application in both basic and translational research but also introduces an approach that unlocks vast possibilities for a deeper comprehension of sarcoma biology.

## MATERIALS AND METHODS

### Tissue culture and GEM model preservation

Human Embryonic Kidney HEK293T cells (RRID:CVCL_0063), human alveolar RMS cell line Rh30 (CVCL_0041) and murine neuroblastoma cells N2A (CCL-131) were purchased from the American Type Culture Collection (ATCC) and maintained in DMEM (Gibco) supplemented with 10% Fetal Bovine Serum and 1% Penicillin/Streptomycin (P/S). IMT_NTRK1/INF_R_153 carrying ETV6-NTRK3 was generated from a primary tumor biopsy obtained from an inflammatory myofibroblastic tumor (IMT) enrolled in the INFORM registry study and cultured in RPMI (Gibco) + 10% FCS + 1% MEM (minimal essential amino acids) + 1% P/S.

For tumor cell purification of mouse sarcoma EPO-GEMMs, existing protocols for primary human sarcomas (Peterziel et al., 2022) were adapted to mouse specimens. GEMM tumors were thoroughly minced with scissors, taken up in 40ml of FCS-free DMEM supplemented with 480 μl of 10 mg/ml Trypsin (Sigma, T9935) and 800 μl of 50 mg/ml Collagenase II (Thermo Fisher, 17101015) and subjected to digestion in a water bath at 37°C for one hour under repeated swirling. If tumor fragments were also desired for allografting, minced tumor fragments were resuspended in FCS-free DMEM and filtered through 200 μm (PluriSelect, 43-50200-50) and 50 μm (PluriSelect, 43-50050-50) cell strainers arranged on a 50 ml falcon tube. This process isolated tumor cell fragments ranging from 50-200 μm, which were then washed into an additional falcon and stored at 4°C until allografting or freezing. To halt enzymatic digestion, 240 μl of 10 mg/ml Trypsin inhibitor (Sigma, T6522) was added. Next, extracellular DNA was digested by a 1:1 mix of 2mg/ml DNase (Sigma, D4527) and 1M magnesium chloride (Fisher Scientific, 15493679) added in 1 to 4 steps of 60 μl each until viscosity decreased and tumor fragments settled. The solution was filtered through a 40 μm cell strainer (Corning, 352340), centrifuged at 400g for 5 minutes, and then resuspended in 2ml ACK lysis buffer (Thermo Fisher, A1049201) for 2-5 minutes for red blood cell lysis. After two washes with 10 ml of FCS-free DMEM, cells were counted (Countess 3 cell counter, Invitrogen, AMQAF2000) and resuspended for seeding or re-engraftment in suitable media. About 0.5 to 1*10^6^ were seeded in 2ml of full DMEM (2D culture) or a spheroid TSM-complete for 3D culture (Peterziel et al., 2022) into 6 well plates (Greiner, 657160 for adherent culture and Thermo Fisher, 174932 for suspension culture). Collagen coating was employed for 2D culture to facilitate attachment for the first passage. After six *in vitro* passages, cell lines were considered established. Dissociation of cells was performed with Trypsin (Sigma, T4049) for 2D cultures and TrypLE (Invitrogen, 12604-013) for 3D cultures. For cryopreservation, about 1-2 million cells, or 1000-3000 tumor spheroids or fragments, were suspended in Synth-a-Freeze Cryopreservation Medium (Gibco, A1254201), placed in gradual freezing aid (Thermo Fisher, #5100-0001), and moved to -80°C before transferal to liquid nitrogen for long-term storage. All *in vitro* lines were cultured in a humidified incubator at 37°C with 5% CO2. The MycoAlert Kit (Biozym 883103) was regularly employed to verify the absence of bacterial contamination with mycoplasma, in all cell lines.

### Plasmids and vectors

Constructs cloned for and used in this study are listed in **Table S4**. For most electroporation experiments, pSB_EF1a_MCS or pPB_EF1a_MCS vectors were used to insert oncogenic open reading frames amplified from human cDNA or ordered as gene strands based on publicly available coding sequences. Assembly was achieved by restriction insertion cloning or Gibson assembly using the NEB Hifi kit (E2621L). For luciferase vectors, a point-mutated version reported to be less immunogenic, albeit slightly less bright was used (Gürlevik et al., 2016). Whole-plasmid sequencing was employed to ensure that plasmids maintained the correct sequences. *KRAS* expression was validated in HEK293T cells via Western Blot after transfection with PEI (Thermo Fisher, BMS1003-A). The ASPS model was generated in a collaborative project with Priya Chudasama on “Immunogenomics characterization of alveolar soft part sarcoma”.

sgRNAs were ordered as oligonucleotides from Sigma-Aldrich and inserted into the PX330 CRISPR vector (Addgene 42230) by digestion with BbsI. Correct insertion was validated by Sanger Sequencing. sgRNAs were designed using Benchling.com, aiming to target the first common exon across all isoforms with highest on- and lowest off-target scores. Five guides per target gene were tested *in vitro* for their editing efficiency in N2A cells after transfection using Lipofectamine 3000 (Thermo Fisher, L3000001). Where available previously published guides were included. sgRNA sequences are listed in **Table S5**.

For *in vivo* electroporation, vectors were amplified by endotoxin-free Giga prep (Qiagen 12391 or Zymo Research D4204). NEB Stable chemically competent E. coli (NEB C3040H) were used and cultured at 32°C. Giga preps were started from validated glycerol stocks in 6ml LB cultures, cultured overnight and used to inoculate 2.5l cultures, harvested the following day for preparation. Final DNA pellets were taken up in about 200µl of PBS and diluted to about 5-10µg/µl. For some experiments the PBS was adjusted to a final concentration of 6 mg/ml of poly-L-glutamate (Sigma- Aldrich, P4761). Plasmids were stored at 4°C for short-term and -20°C for long-term storage. For electroporation, 4µg of transposase was mixed with 8µg of each transposon vector, 8µg of each CRISPR vector and 8µg of each reporter vector as outlined in results. For P30 EPO the plasmid mixture was adjusted to 1:100 with methylene blue dye (Sigma 50484) in PBS in a final volume of 25µl per leg. For P0 EPO, methylene blue was used at 1:25 with a final volume of 5µl. For Sleeping Beauty vectors (SB) SB13 transposase was used as *in vitro* experiments did not show superiority of SB100X over SB13. For PiggyBac (PB) vectors, PiggyBac transposase was used.

### Western blotting

Cells were harvested, washed with PBS, and resuspended in RIPA buffer (Cell Signaling, 8906) with protease inhibitors (Sigma-Aldrich, 11836170001) for 30 minutes on ice under regular vortexing. After centrifugation (15 minutes at 17,000 g, 4°C), protein lysates were quantified using the BCA protein assay (Thermo Fisher, 23227). Samples were adjusted, denatured in 2X Laemmli at 95°C for 5 minutes, and loaded onto 4-15% protein gels (Biorad, Biorad 456-1084) at 30μg per well. Semi-dry blotting (Biorad, #1704150) transferred proteins to a PVDF membrane, pre-activated with methanol. The membrane was blocked, incubated with primary antibody overnight, washed, and incubated with a secondary antibody. After a final wash, the membrane was incubated with ECL solution (Perkin-Elmer, NEL103001EA) and developed on an Amersham Imager 680. For β-actin antibody (already HRP-coupled), the secondary antibody step could be omitted. A list of utilized antibodies is delineated in **Table S6**.

### Animal studies

All animal experiments conducted in this study were carefully planned and approved by the government of North Baden as the responsible legal authority (G-36/19, G-2/20, G-3/20). The study adhered to the ARRIVE guidelines, European Community and GV-SOLAS recommendations (86/609/EEC), and United Kingdom Coordinating Committee on Cancer Research (UKCCCR) guidelines for the welfare and use of animals in cancer research. Conscientious application of the 3R guideline (replacement, reduction, refinement) was emphasized, prioritizing the reduction of potential suffering for the animals. Animals for this study were purchased from Janvier laboratories and housed at the central DKFZ animal facility under Specific Opportunist Pathogen-Free (SOPF) conditions, utilizing individually ventilated cages. Animals had *ad libitum* access to food and water. Daily assessments of their well-being were carried out by certified animal caretakers. Details on utilized mouse strains (CD-1 and C57BL6/J) are given in **Table S7**.

### *In vivo* electroporation

Electroporation of P30 (4-6 weeks old) animals: Preemptive oral analgesia (4 mg/ml Metamizol in drinking water *ad libitum*, sweetened to achieve a final concentration of 1.5% glucose) commenced one day before surgery and persisted for three days thereafter. Additionally, animals received a single dose of 200 mg/kg metamizole subcutaneously under anesthesia. Anesthesia was induced with 1.5-2.5% isoflurane under close monitoring of vital functions before shaving the legs, disinfecting the operation field, and initiating the procedure. A warming mat maintained stable body temperature. Bepanthen ointment protected eyes from drying or keratitis. Mice were positioned on their backs on a sterile drape with limbs gently secured. The quadriceps femoris muscle was surgically exposed bilaterally. For most experiments, 25μl of hyaluronidase was intramuscularly injected to enhance transfection efficiency before temporarily closing the skin with wound clips. After a two-hour rest, clips were removed under anesthesia. In experiments without hyaluronidase pre-treatment, muscle exposure was directly followed by electroporation. The blue plasmid mixture was injected into the thigh muscle perpendicularly to the muscle fiber direction under visual control using a calibrated Hamilton glass syringe (25 μl, Model 702 RN, CAL7636-01) with a 28G needle (Removable needle, 28G, point style 4, 7803-02). Thereafter, the exposed muscle was directly placed between two 5 mm platinum plate electrodes (Nepagene, CUY650P0.5-3), and 5 unidirectional 100V pulses of 35 ms length with 500 ms intervals (or other conditions as outlined in results) were applied using a calibrated electroporator (Nepagene, NEPA21). The procedure concluded with a final disinfection of the operation field and wound closure through continuous suturing before the animals were placed in a separate cage to recover under close supervision. The procedure was well-tolerated.

Electroporation of P0 (newborn) animals: Mother animals received additional nesting material approximately one week before birth to mitigate the risk of offspring rejection post-procedure. Following birth, pups underwent a brief separation of approximately 10-15 minutes from the dam, during which EPO was conducted under anesthesia with 1.5-2.5% isoflurane. Pups were collectively placed on a warming mat beneath a sterile drape to maintain stable body temperature. For EPO, mice were positioned on their backs, and limbs were gently secured with sterile tape after gentle disinfecting the operation field. Due to the delicate nature of newborn skin, surgical exposure of the muscle was omitted. Instead, 5 μl of plasmid mix, with 1:25 diluted methylene blue, was transcutaneously injected into the thigh muscle of both legs using a calibrated Hamilton glass syringe (5 μl, Model 75 RN, CAL7634- 01) with a 30G needle (Removable needle, point style 4, 7803-07). The higher concentration of methylene blue allowed visually controlled injection, albeit slightly less precise than in P30 mice. Subsequently, the leg was placed between two 5mm platinum plate electrodes, and 5 unidirectional 70 V pulses of 35 ms length with 500 ms intervals (or other conditions as outlined in results) were applied using the Nepa21 electroporator. Post EPO, mouse pups were reunited with their mother and left undisturbed. The procedure was well-tolerated. All offspring from both CD1 and C57BL6/J mice were consistently well-accepted.

### Tumor surveillance in *vivo*

Caliper measurements: In addition to daily health assessments by certified animal caretakers, comprehensive weekly examinations were carried out by scientific staff to detect tumors through thigh palpation and identify associated signs of disease. Regular weight measurements were also performed. Once a palpable tumor emerged, its length and width were consistently measured using a digital caliper (Fine Science Tools, 30087-00), typically 3 times per week, with daily monitoring for rapidly growing tumors. Tumor volume was determined using the formula: (length * width^2^)/2. Animals were observed for up to one year post-electroporation unless tumor growth reached a maximum of 15mm in one dimension or other termination criteria were met, which included weight loss of up to 20%, apathy, abnormal posture, piloerection, respiratory problems, and specific signs on the Grimace Scale (constricted eyelids, sunken eyes, flattened ears) (Whittaker et al., 2021), invasive growth into thigh muscles causing functional limitations (lameness), disability, or pain, exulcerations or automutilations. Neonatal mice were terminated if they exhibited the absence of a milk spot, cannibalism, rejection by the mother, color change (from pink to blue or pale), or lack of locomotion in response to touch stimuli. Mice were killed by cervical dislocation with or without deep narcosis or increasing CO_2_ concentrations. Newborns until P5 were killed by decapitation.

*In vivo* bioluminescence imaging: When utilizing luciferase as a reporter gene, regular monitoring of gene expression, transfection efficiency, and tumor growth was conducted using IVIS (*In vivo* imaging system) bioluminescence imaging. D-luciferin (Enzo, 45784443) was prepared in PBS (15 mg/ml), sterile-filtered, aliquoted into light-protected vials, and stored at -20°C until use. Mice were anesthetized with 1.5-2.5% isoflurane, injected intraperitoneally (i.p.) with luciferin (10 μl/g of mouse weight), and positioned in the IVIS chamber on their backs with isoflurane administered through a mouthpiece. Bepanthen ointment was used to protect their eyes. A 10-minute incubation period was followed by 7 minutes of image acquisition, determined to be optimal during the plateau phase of the IVIS signal based on pilot experiments. Typically, three animals were imaged simultaneously. When four to five animals were imaged concurrently, an XFOV-24 lens was used. After imaging, mice were placed in a separate cage to recover. Living Image software (PerkinElmer, version 4.5.5) was used for analysis. Regions of interest (ROIs) were defined around the electroporated regions to quantify IVIS signal intensity as photons/second.

### Syngeneic allografting of mouse tumors

Primary and *in vitro* cultured murine tumor material from EPO-GEMMs was purified and dissociated as outlined above (cell culture section), quantified and re-engrafted syngeneically into wildtype C57BL6/J recipient mice. Dissociated cells were counted using an automated cell counter (Countess 3, Invitrogen, AMQAF2000), while 50-200µm sized and tumor fragments and spheroids were manually counted in 10*5 μl droplets on a petri dish under a light microscope (Zeiss, 491237-9880-010). The mean count of five droplets was used to estimate the total number of fragments or spheroids. Cells, spheroids, and fragments were then centrifuged and resuspended at concentrations of 1*10^6^ primary or cultured cells and 1000 spheroids or fragments per 12.5µl of FCS-free DMEM, which were kept on ice until injection. Perioperative preparation was performed analogously to P30 EPO, except for shaving, which was exclusively performed on the left leg. Mice were placed on their backs, and extremities were gently fixed with sterile tape. A sterile pen was positioned diagonally beneath the left leg to expose the thigh muscle. Just before engraftment, solutions containing tumor fragments, tumor cells, or tumor spheroids were resuspended ice in 1:1 ratio with Matrigel (Corning, 354277 or Thermo Fisher, A1413202) and loaded into a calibrated Hamilton glass syringe (25 μl, Model 702 RN, CAL7636-01) equipped with a 28G needle (Removable needle, 28G, point style 4, #7803-02). The injection was performed transcutaneously into the thigh muscle perpendicularly to the muscle fiber direction under visual control. Subsequently, mice were placed in a separate cage to recover.

### Genetic manipulation of murine tumor lines *in vitro*

Murine tumor lines were equipped with human B7-H3 target antigen (ENST00000318443.10) or human CD19 antigen (<u>ENST00000538922.8</u>) by lentiviral transduction and puromycin selection using standard protocols. In short, human CD276 or CD19 was amplified from cDNA and truncated to contain only the first 10-15 intracellular amino acids. The PCR product was cloned via HiFi DNA Assembly into the pCDH lentiviral backbone to form pCDH-hCD276t/hCD19t-P2A-Pac-T2A-FlucL272A. Murine SAM cell lines were transduced using standard lentivirus practice and continuously selected with 2 µg/ml puromycin.

### Generation of murine CAR-T cells

Murine T lymphocytes were isolated from the spleens of 8-12 week-old wildtype C57BL/6J mice using the Pan T Cell Isolation Kit II, mouse (Miltenyi Biotec, 130-095-130) following the manufacturer’s instructions. After isolation, cells were cultured in complete TexMACS medium (Miltenyi Biotec, 130-097-196) supplemented with 10 % FBS, 1 % PenStrep, 50 µM β-Mercaptoethanol (Gibco, 31350010) and 50 IU/ml premium-grade human IL2 (Miltenyi Biotec, 130-097-746) at 1x10^6^ cells per mL and cm^2^. Cells were stimulated using the T Cell Activation/Expansion Kit, mouse (Miltenyi Biotec, 130-097-627) with a 2:1 bead:cell ratio.

The chimeric antigen receptor against human CD276 (Du et al., 2019) and CD19 (Maude et al., 2018)together with the Thy1.1 surface antigen (Addgene 154194)(Umkehrer et al., 2021) was cloned into pMSCV-U6sgRNA(BbsI)-PGKpuro2ABFP (Addgene 102796)(Henriksson et al., 2019) in place of the puro2ABFP sequences, resulting in pMSCV-U6-sgRNA(BbsI)- pgk-CD276_CAR-P2A-Thy1.1 and and pMSCV-U6-sgRNA(BbsI)-pgk-CD19_CAR-P2A- Thy1.1.

Ecotropic retrovirus was produced using the Platinum Eco cell line (Morita et al., 2000) (Biocat, RV-101-GVO-CB). Briefly, cells were harvested and seeded with 10.5 x10^6^ cells per 15-cm-dish in complete DMEM without Pen/Strep 24 h prior to transfection. Cells were transfected using Turbofect (Thermo Scientific, R0532) with 40 µg plasmid. Medium was changed after 12-18 h post transfection and cells were incubated for another 24-30 h before virus harvest. Virus was harvested, cleared once by centrifugation (800 g, 10 min, 32 °C), pelleted overnight using Retro-X-Concentrator (Takara Bio, 631456) and resuspended in complete TexMACS the next morning. Meanwhile, untreated 6-well cell culture plates (Falcon, 351146) were coated with 15 µg/ml retronectin solution (Takara Bio, T100A) overnight at 4 °C. 24 h after T cell stimulation, T cells were pelleted and resuspended in complete TexMACS. Per one retronectin-coated 6-well, 3.3 ×10^6^ T cells were resuspended in 2 mL medium, and 2 ml concentrated virus supernatant (equaling virus yield from one 10- cm-dish) was added. Cells were then spin-infected by centrifugation at 2,000 g for 90 min at 32 °C and incubated for another 12-24 h before transfer to 12 well plates at 2 ×10^6^ cells per well 2 ml of fresh medium. At day 5 cells were transferred to larger culture flasks for further expansion. Experiments were performed at day 10. Control T cells underwent the same procedure in the absence of any virus.

### Flow Cytometry

Human B7-H3 expression on murine tumor cells and expression of immune cell markers CD3 and Anti-B7-H3-CAR markers CD90.1 and myc-tag on murine T cells were quantified using FACS. For B7-H3 FACS, tumor cells were harvested by incubation with PBS and EDTA (20mM). Cells were stained with ZombieViolet (Biolegend, 423113, 1:500) and anti- CD276-FITC (Miltenyi Biotec, 130-118-697, 1:50) following the manufacturer’s recommendations. CAR-T cells were stained using anti-CD90.1-VioBlue (Miltenyi Biotec, 130-112-878, 1:50), anti-c-myc-biotin and anti-biotin-PE (Miltenyi Biotec, 130-124-899 and 130-111-068, 1:50 each) and Helix NP NIR (Biolegend, 425301) following the manufacturer’s instructions. Fc block was performed using FcR blocking reagent (Miltenyi Biotec, 130-092-575). Staining buffer consisted of PBS with 2 mM EDTA and 0.5% BSA, except for ZombieViolet where PBS with EDTA was used. All samples were measured on a LSR Fortessa (BD Bioscience), compensated on the cytometer using single-stain controls or beads (Miltenyi Biotec, 130-104-693; and Biolegend, 424602) and analyzed in FlowJo v10 (BD Life Science). For analysis, cells were gated based on FSC/SSC signals and only live cells were used for marker analysis.

### Preclinical treatment studies

NTRK inhibitor treatment *in vitro*: 500-1000 cells per well cultured in full DMEM were seeded into black flat-bottom 384 well plates (Greiner, 781091) for each cell line based on preliminary experiments to determine the appropriate cell numbers based on baseline growth rates. Each drug was pre-printed into 384 well plates in 10 doses, ranging from 0 to 10,000 nM of Larotrectinib (Medchem Express, HY-12866) and Repotrectinib (Medchem Express, HY-103022) taken up in 99.9% DMSO (Sigma, D8418) in semi-logarithmic increments (in amounts of 0, 1, 3.16, 10, 31.6, 100, 316, 1000, 3160, 10000 nM). 100 µM of Benzethonium chloride (Sigma, PHR1425) was used as a positive control. Staurosporin (TargetMol, T6680) as a dose-dependent therapy response control with one technical replicate per dose and cell line (in amounts of 0.1, 1, 10, 100, 1000 nM). For all other groups, four technical replicates were used per dose and drug for each cell line. Edges of the plates were filled with PBS to avoid edge effects. Order of drugs and doses was based on a random distribution to avoid batch effects. Treatment effects were quantified 72 hours after cell seeding using calorimetric ATP measurement with CellTiterGlo 2.0 (Promega, G7572) according to the manufacturer’s protocol and as described previously (Peterziel et al., 2022) using an Infinite M Plex plate reader (Tecan). Data was analyzed with using the previously described iTrex algorithm (ElHarouni et al., 2022) for quality control and to condense dose-response curves into representative drug sensitivity scores (DSS) ranging from 0-50, 0 indicating resistance, >10 indicating sensitivity.

CAR-T treatment *in vitro*: 10^4^ target cells (SA797_tll_2D-hCD276t) were seeded in 96-well plates (Greiner, 655098) 24 h prior to addition of T cells. After 24 h of incubation, control T- or anti-CD276-CAR-T cells were added to each well with indicated ratios relative to the initially seeded tumor cell numbers. D-luciferin (Abcam, 143655) was added to a final concentration of 150 µg/ml. Luminescence was measured every 24 h using an Infinite M Plex plate reader (Tecan) with 1 second integration time. Blank values were subtracted from all other values, and percentage of killed cells was calculated as (1-(mean treated wells / mean control wells)).

CAR-T treatment *in vivo*: Tumors were induced by engrafting 150,000 murine aRMS previously transduced with human B7-H3 with 1:1 mixture of matrigel (Corning, 356234) into the left thigh muscle of 7-week old female C57BL/6J mice. Treatment was started 10 days post tumor cell engraftment using *in vitro* expanded T cells at day 8 of expansion. Activation beads were removed using the MACSiMag magnet (Miltenyi Biotec, 130-092- 168) and cells were taken up in PBS. Cell density was adjusted to indicated numbers based on live cells and corrected for the fraction of CAR-positive T cells in 100 µl injection volume, applied via tail vein injection. Tumor-bearing mice were randomized for treatment.

### Tissue removal and processing

Euthanized mice were carefully examined for metastatic disease of internal organs after removing the skin. Tumors were photographed. Some animals underwent macroscopic examination of GFP expression using a fluorescent dissection microscope (Leica, 25716). Tumors were surgically separated from the femur and surrounding organs. To prevent cross- contamination between tumors, all instruments were meticulously disinfected before handling a new tumor. The tumor was dissected into multiple sections on an inverted Petri dish using a microtome blade (VWR, 720-2369). One cross-section was placed in a histo cassette (Sigma, H0792-1CS) and fixed in formalin for 2-3 days before transfer to 50% (v/v) ethanol at 4°C until further processing. Another cross-section was embedded in cryo-embedding resin (OCT compound, Tissue-Tek, 14291) and placed in a cryomold (Tissue-Tek, 14292), which was set on a metal rack over dry ice for homogeneous snap freezing before transfer to -80°C. The remaining tissue was sectioned into small tumor pieces (20-30mg), with portions allocated for nucleic acid purification for molecular analysis (snap-frozen on dry ice) and tumor cell isolation for cell culture and allografting into recipient mice.

### Nucleic acid extraction

Before nucleic acid extraction, all surfaces and instruments underwent cleaning with RNase decontamination reagent (Thermo Fisher, 7000TS1). DNA and RNA were extracted from the same tumor tissue piece, approximately 20-30 mg in weight, using Qiagen’s DNA/RNA AllPrep kit (80204) following the manufacturer’s protocol. If only DNA was required, the Qiagen DNeasy kit (69506) was employed for purification. Tissue homogenization was accomplished using DNAse/RNAse-free pestles (Carl Roth, CXH7.1), swirled in 1.5ml Eppendorf tubes with a cordless pestle motor (DWK Life Sciences, 749540-0000). Lysates were subsequently passed through a QIAshredder column (Qiagen, 79654) to eliminate debris before proceeding with nucleic acid purification. Purified DNA and RNA were maintained on ice, concentration was determined using Nanodrop and Qubit, and then stored at -20°C (DNA) and -80°C (RNA) for subsequent analyses.

### Oncogene PCR from genomic DNA

Genomic DNA, extracted from tumor samples and plasmid DNA as positive controls, served as the template for polymerase chain reaction (PCR). 1 µl of genomic or plasmid DNA, containing 10-100 ng of DNA, was combined with 12.5 µl of 2X Red HS Mastermix (Biozym, #331126L), 1.25 µl of the forward primer, 1.25 µl of the reverse primer, and 9 µl of water. PCR products were loaded onto 0.7-1% (w/v) agarose gels, along with 1 kb (NEB, N3232L) and 100 bp (NEB, N3231) DNA ladders. Electrophoresis was conducted at 80- 110V until satisfactory separation was achieved, followed by imaging using a Gel doc imager (Biorad, 170-8170). Primer sequences and PCR conditions for genotyping PCRs are outlined in **Table S8**.

### Indel analysis from genomic DNA

To assess the efficacy of CRISPR-mediated editing of tumor suppressor genes, primers were designed to amplify the corresponding genomic loci spanning approximately 500-700 base pairs around the sgRNA target site. PCR products were purified using NucleoSpin Gel and PCR Clean-up Kit (MACHEREY-NAGEL, 740.609.250) and subsequently subjected to Sanger sequencing through Microsynth Seqlab services, utilizing the corresponding forward primer. The obtained sequences were aligned with the wildtype sequences derived from mouse tail genomic DNA, using the TIDE algorithm (Tracking of Indels by Decomposition) (Brinkman et al., 2014) (http://shinyapps.datacurators.nl/tide/) to calculate the percentage of insertions and deletions. Primer sequences and PCR conditions for TIDE PCRs are outlined in **Table S5**.

### Immunohistochemistry and mulitplexed immunofluorescence imaging

Immunohistochemistry: After fixation in 10% (v/v) buffered formalin (Sigma, HT501128) for 2-3 days, tissue sections underwent a stepwise rehydration process in an alcohol series using an automated tissue processor (Leica, ASP300S) until reaching a concentration of ≥99% (v/v) ethanol. Subsequently, the tissue cassettes were transferred to intermediate xylene and embedded in paraffin (Leica, 14039357258). 3-4 μm paraffin sections were prepared with a HM 355S microtome (Fisher Scientific, 10862110), deparaffinized and rehydrated up to 96% ethanol (v/v). H&E and PAS stainings were performed with standard protocols. For H&E, 5 minutes of Haemalaun nuclear staining (Carl Roth, T865.3) was followed by 5 minutes of rinsing with water and 20-30 seconds of Eosin solution (Merck, 115935, 100ml supplemented with one drop of acetic acid, Merck, 1.00063). For PAS staining, periodic acid (Merck, 100524) was applied for 10 minutes, Schiff’s reagent (Merck, 1.09033) for 5 minutes, Haemalaun nuclear staining (Carl Roth, T865.3) for 1 minute, and 5 minutes of rinsing. For immunohistochemistry, primary antibodies were typically diluted in antibody Diluent (DAKO, S2022) with 2% milk (v/v) for 90 minutes at 37°C, followed by four washes with TBST-T and application of biotinylated secondary antibodies against the species of the primary antibody, typically diluted 1:500 in TBS-T for 30 minutes at 37°C.

This was followed by three washes and treatment with 1:200 alkaline phosphatase streptavidin (Vector, SA-5100) in TBS-T for 30 minutes at 37°C. Signal amplification, if necessary, involved another round of secondary antibody and alkaline phosphatase streptavidin treatment. Finally, substrate red (Dako, K5005) was applied for 10 minutes at room temperature before slides were washed, counterstained with Haemalaun (Carl Roth, T865.3), and mounted with Aquatex (Sigma, 1.08562.0050). Antibodies and corresponding antigen retrieval methods are listed in **Table S6**. Slides were scanned using the Aperio AT2 slide scanner (Leica, 23AT2100) at 40x magnification. Morphological assessment and quantification of immunohistochemistry and morphological tumor features were conducted with the expertise of pathologist Felix Kommoss in blinded fashion. Digitized image files were analyzed using QuPath software (version 0.4.1) and assembled into final figures using Affinity Designer software (version 1.10.5).

Multiplexed immunofluorescence imaging: Cryo-embedded mouse tumors (OCT compound, Tissue-Tek, #14291) were cut to 4-5µm with a cryostat (Leica CM 1950, Leica Biosystems) and positioned onto SuperFrost plus slides (R.Langenbrinck, 03-0060). Slides were stored at - 80 °C until use. Tissue fixation was performed with 4 % paraformaldehyde (Thermo Scientific, J19943.K2) for 10 minutes at room temperature. Subsequently, slides were mounted onto the respective MACSwell imaging frame (Miltenyi Biotec, 130-124-673), washed 3 times with MACSima™ Running Buffer (Miltenyi Biotec, 130-121-565) before nuclear staining was performed with DAPI (Miltenyi Biotec, 130-111-570) diluted 1:10 in running buffer. Slides were washed three times with running buffer and submitted to MICS (MACSima^TM^ imaging cyclic staining). Antibodies were diluted with Running Buffer in a MACSwell™ Deepwell Plates (Miltenyi Biotec, 130-126-865) in a total volume of 1050 µl. Spatial analysis using MICS comprised fully automated iterative cycles of ultra-high content immunofluorescence based on fluorochrome-labelled antibody staining, image acquisition, and fluorochrome removal. Images were generated according to the manufacturer’s instructions and as described before (Kinkhabwala et al., 2022). In brief, regions of interest were defined based on the DAPI signal and the focus was set using hardware autofocus settings. Raw data was processed with the MACS® iQ View image analysis software (Version 1.2.2) as described before (Kinkhabwala et al., 2022). Processing included automated optimal exposure time selection, calibration correction, stitching of fields of view (FoVs), and subtraction of pre-stain bleach images. Image quality control was assessed, data was analyzed, and MICS data was visualized using MACS® iQ. Utilized antibodies and dilutions are detailed in **Table S6**.

### DNA methylation and CNV analysis of murine tumors and tissue

Genome-wide methylation profiling was performed using the Illumina Infinium Mouse Methylation BeadChip covering > 285,000 CpG sites distributed over the mouse genome. A minimum of 500ng of genomic DNA extracted from frozen tissue was submitted per sample. DNA quality was ensured by digital gel electrophoresis (Agilent, G2939BA). IDAT files were obtained and processed using the R package sesame, version 1.16.1 (SEnsible Step-wise Analysis of DNA MEthylation BeadChips)(Zhou et al., 2018) to generate normalized beta values. Probe intensities underwent background correction using the p-value with the out-of- band array hybridization approach, followed by a normal-exponential out-of-band approach. Dye bias correction was performed by aligning green and red to the midpoint using the dyeBiasCorrTypeINorm method in sesame. Probes targeting the X and Y chromosomes were excluded. Clustering analysis was performed with R packages Rtsne and umap based on the 2,000 or 10,000 most variably methylated probes, with perplexity values set to 5 for tSNE and 15 for umap clustering. To infer CNV profiles, the method described in R package conumee, version 1.32.0, was adapted for the mouse array. Specifically, a panel of n=60 normal tissue idat files from C57BL6 mice, kindly provided by Marc Zuckermann and Tuyu Zheng (DKFZ) underwent the same sesame correction pipeline, and total probe intensities were quantified across all probes in tumor and normal samples. The background ratio of cancer sample to normal control intensities was determined using the slope of a linear model. Subsequently, the log base 2 ratio of observed vs. expected intensity was calculated for every probe. Probes were binned using sesame according to their mm285 array manifest, utilizing the getBinCoordinates function. For heatmap visualization of copy number variation, the color range representing the log fold change in probe intensities was set to -1 to +1.

### RNA sequencing analysis of murine tumors and tissue

RNA exclusively sourced from fresh frozen tissues was, quality-controlled by digital gel electrophoresis (Agilent, G2939BA). Only samples with RNA integrity scores (RIN) exceeding 7 proceeded to library preparation using the TruSeq Stranded total mRNA protocol, starting with 50 μl of undegraded RNA at concentrations ranging from 50 to 80 ng/μl per sample. Sequencing was carried out on Illumina’s NovaSeq 6000 S1 or S4 flow cell with paired-end 50 bp reads, yielding an average of approximately 20 million reads per sample. Sequencing reads were aligned to the mouse reference genome (GRCm38mm10) by DKFZ’s Omics IT and Data Management Core Facility (ODCF) using the One Touch Pipeline (OTP)264 / RNAseq workflow pipeline, version 1.3.0 (STAR Version 2.5.3a, Merging/duplication marking program: Sambamba Version 0.6.5. SAMtools program: Version 1.6.), resulting in raw counts, RPKM, and TPM values. Initial exploratory data analysis utilized the iDEP web collection of R packages (Ge et al., 2018). Further differential gene expression analysis was conducted with DeSeq2 in R Studio, version 2022.07.1. Additional R packages such as fgsea, clusterProfiler, biomaRt, and org.Mm.eg.db were employed for Gene Set Enrichment Analysis (GSEA) using publicly available gene set collections downloaded from MSigDB (Molecular Signatures Database). For cell deconvolution, RNA Seq data was processed using CibersortX in accordance with the developer’s manual (https://cibersortx.stanford.edu)(Newman et al., 2019). The ’Tabula muris’ dataset, a publicly accessible collection of single-cell RNA sequencing data from mice, including >100,000 annotated single cells representing more than 130 cell types across 20 different organs (Tabula Muris Consortium et al., 2018) was used to create a reference matrix for cell type annotation. Batch correction parameters were set to S mode (recommended for single cell reference data).

### Cross-species transcriptome analysis of mouse EPO-GEMMs and human sarcomas

To compare RNA-sequencing profiles between mouse and human sarcoma specimens, data of 2222 human sarcomas (TCGA, St. Judes and INFORM), representing more than 20 common and rare soft-tissue and bone sarcoma entities, was assessed for the top 500 most differentially expressed genes in DEseq2 v1.30.1 (log_2_FC >2, FDR< 0.05) in each subtype vs all other subtypes. The top 500 most differentially expressed genes were determined in the same way for n=63 mouse samples of primary GEMMs representing 10 subtypes entities. All selected human genes were matched to orthologs in mouse genes. In cases with no orthologs these genes could not be considered in the comparative analysis. The lists of mouse genes and orthologs from human genes were combined to a total of 2636 genes. This gene list was subsequently used to create matrix and batch correction was performed for species and sample type (“Muscle control” and dataset (St. Jude, TCGA and INFORM) using Harmony v0.10 (Korsunsky et al., 2019), implemented in R v4.0.3. These batch corrected values were then subjected to t-SNE clustering, excluding non-relevant entities that could not be clearly assigned to one entity or did not have matching mouse samples (e.g. Liposarcoma, Osteosarcoma or EwS).

### Statistical analysis

Statistical analysis was performed as outlined in the respective figure captions for each experiment using R v4.0.3. Wherever suitable the Bonferroni-Holm method was used to correct for multiple testing. The provided sample size (n) indicates biological replicates. Group sizes for *in vivo* experiments were determined through statistical consultation in the Department of Biostatistics at the DKFZ, which included in silico simulations. Group allocation for preclinical treatment studies was determined through random distribution. Outcome assessment for preclinical treatment trials were done in blinded fashion. Other group allocations and outcome assessments were performed in non-blinded fashion. Survival was measured using the Kaplan-Maier method and log-rank tests. The threshold for significance was set to P < 0.05.

## Figure preparation

Data was plotted using R v4.0.3 and Graphpad Prism version 8.4.3. Affinity Designer version 1.10.5. and Biorender.com were used to generate graphical illustrations and arrange panels into final figures.

## AUTHORS’ CONTRIBUTIONS

R.I. conceived the study, designed, performed, and analyzed the experiments, and wrote the manuscript. A.B. conceived and coordinated the study, designed experiments, acquired funding and wrote the manuscript. S.M.P. acquired resources, funding and writing-review. D.B., S.P., M.H., S.S., C.S., F.H.G., M.v.E., and F.C-A. performed and analyzed some experiments. F.K.F.K., T.G.P.G., and C.V. performed expert histopathology review. E.S.Z., R.A., M.S., and C.B. performed bioinformatic analyses. D.L., L.S., L.W., C.W., and C.Schm. assisted with experiments. I.Oe., H.P., K.J., P.Ch., C. Scho., and P.G. provided essential materials and information. All authors read, revised and approved the final manuscript for publication.

## ACKNOWLEDGEMENTS

We thank Daisuke Kawauchi, Joseph Leibold and Kerstin Dell for insightful discussions regarding the development of a muscle-specific electroporation method and members of the soft-tissue sarcoma lab for feedback and fruitful discussions. Darjus Tschaharganeh, Hsuan-An Chen (Scott Lowe lab) and Rienk Offringa for sharing reagents. We thank the light microscopy core facility, the genomics and proteomics core facility and the small animal imaging center, along with the outstanding and dedicated team of animal caretakers at the DKFZ, for their exceptional support and services. We thank Tim Holland-Letz and the Department of Biostatistics of the DKFZ for statistical consultation. We thank Veronika Eckel and Alexander Brobeil from the Tissue Bank of the National Center for Tumor Diseases (NCT) Heidelberg, Germany for their excellent services in scanning of histology slides. We express our gratitude to Marie Koloseus for her preliminary work in establishing a mouse antibody panel for multiplexed immunofluorescence imaging. We thank Veronika Bahlinger, Christian Schürch and Benjamin Mayer for supplying additional human histology specimens. We express our gratitude to Katharina Hartwig for her valuable contribution to graphic design. This project has received funding from the European Research Council (ERC) under the European Union’s Horizon 2020 research and innovation program (grant agreement n° 805338) (A.B.) as well as the Physician-Scientist Program of the University of Heidelberg, Faculty of Medicine (R.I.). D.B. is supported by the DKFZ International PhD Program. Work in the laboratories of A.B., T.G.P.G., S.M.F., P.C. and C.S. is supported by the HEROES-AYA (Heterogeneity, Evolution, and Resistance in Oncogenic Fusion Gene- Expressing Sarcomas Affecting Adolescents and Young Adults) consortium within the National Decade Against Cancer of the German Federal Ministry of Education and Research (01KD2207). T.G.P.G. acknowledges support of the Barbara and Wilfried Mohr foundation. P.C. receives support from an Emmy Noether Program Grant CH2302 of the DFG (German Research Foundation). Finally, we express our deepest gratitude to all pediatric and adult patients affected by sarcomas who have provided us with inspiration and motivation to pursue this work.

## SUPPLEMENTAL FIGURE LEGENDS

**Figure S1.**
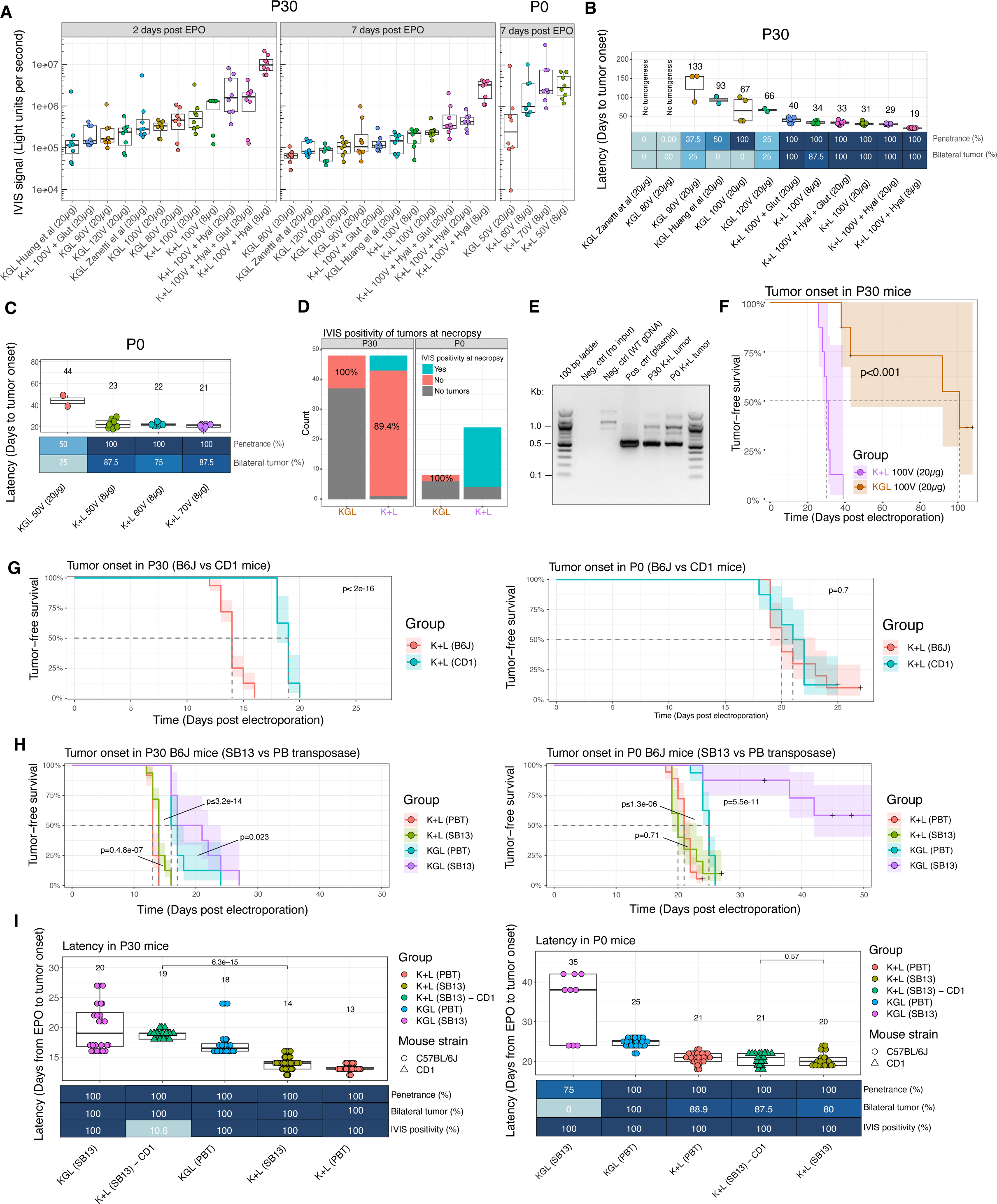
Optimization of transfection efficiency and tumor take in CD1 and C57BL/6J mice **A)** Quantification of IVIS signal at day 2 and 7 after electroporation as a surrogate for muscle transfection efficiency in P30 and P0 CD1 mice (IVIS at day 2 was not feasible in P0 group). Electroporation settings labelled Huang et al (Huang et al., 2017) and Zanetti et (Zaneti et al., 2019) al were obtained from literature. **B-C)** Boxplots of tumor-free survival in CD1 mice ordered by median from high to low with attached heatmaps of penetrance and laterality of tumor onset in P30 (**B**) and P0 (**C**) animals. Mean plotted on top of boxplot. **D)** IVIS positivity of tumors in CD1 mice before necropsy. **E)** Detection of Luciferase transposon in tumor gDNA by PCR. **F)** Tumor-free survival between K+L and KGL tumors in CD1 mice. **G)** Tumor-free survival using K+L vectors in B6J compared to CD1 mice upon P30 and P0 EPO. **H)** Tumor-free survival in B6J mice using K+L vs KGL vectors and SB13 vs PBT transposase upon P30 and P0 EPO. **I)** Boxplots of tumor-free survival in B6J and CD1 mice using the ideal electroporation procedure as determined in panels **A-C**, ordered by median from high to low with attached heatmaps of penetrance, laterality of tumor onset and IVIS positivity before necropsy in P30 and P0 animals. Mean plotted on top of boxplot.

**Figure S2.**
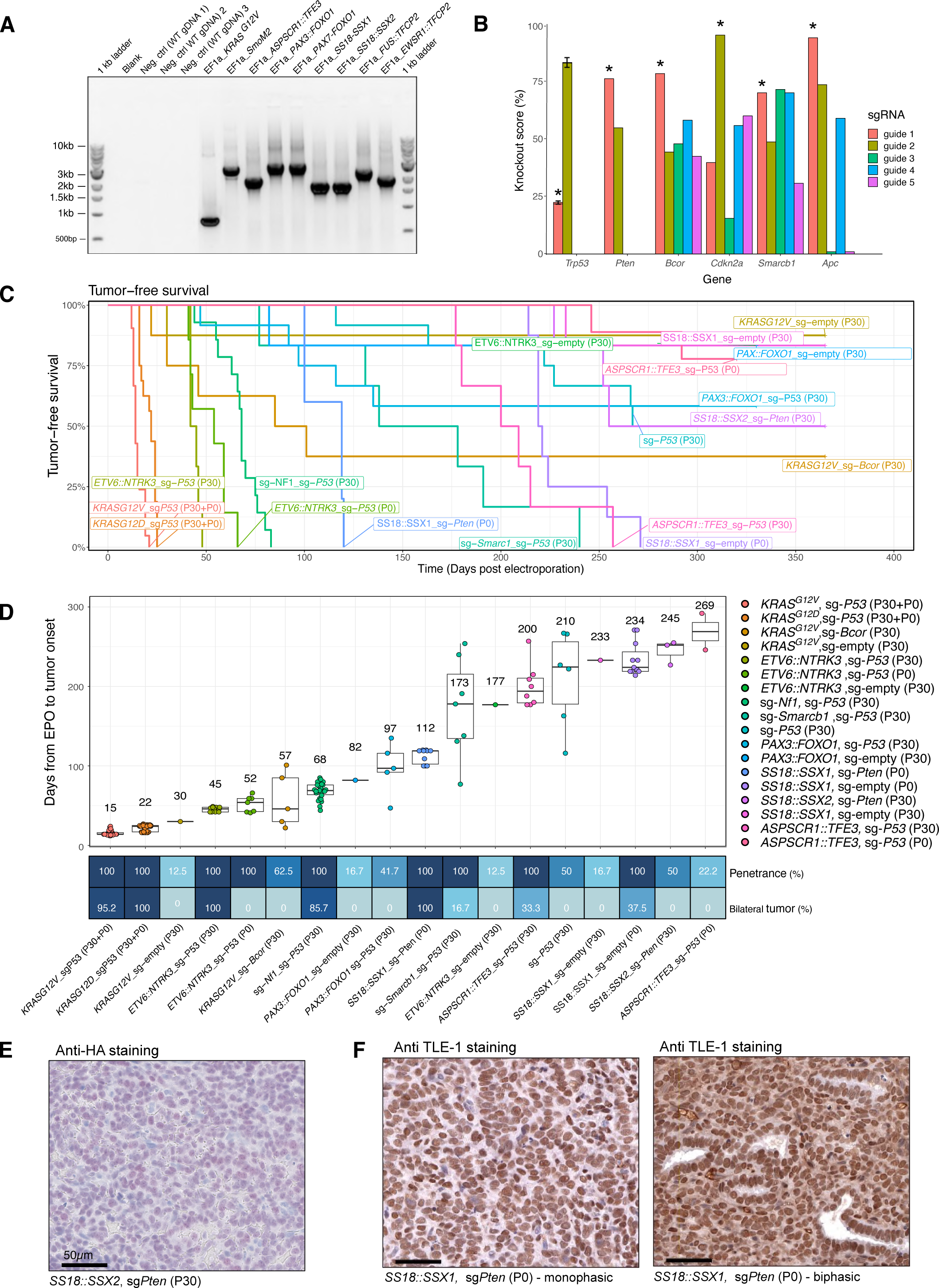
Efficiency of sarcoma induction in mice A) Exemplary genotyping PCRs of transposon vectors used to induce tumors *in vivo*. B) Knockout scores quantified using TIDE analysis in N2A cells (murine neuroblastoma line), three days after transient transfection with PX330 vectors carrying the respective sgRNAs. *Trp53* guide 1 (Dow et al., 2015) shows a falsely low knockout score due to a base mismatch in the *Trp53* target region in N2A cells. Experiments for *Trp53* were performed with n = 3 (SD depicted) to ensure reproducibility of the assay. C) Kaplan-Meier curves of tumor free survival of all genetic combinations that were successful in tumor induction (n=18). D) Tumor-free survival across the mouse sarcoma cohort from panel C as boxplots sorted by median onset with attached heatmaps of penetrance and bilateral tumor fraction. Mean plotted on top of boxplot. E) Anti-HA IHC staining of murine SS, driven by *Flag-HA- SS18::SSX2*. F) Anti TLE-1 IHC staining of murine SS.

**Figure S3.**
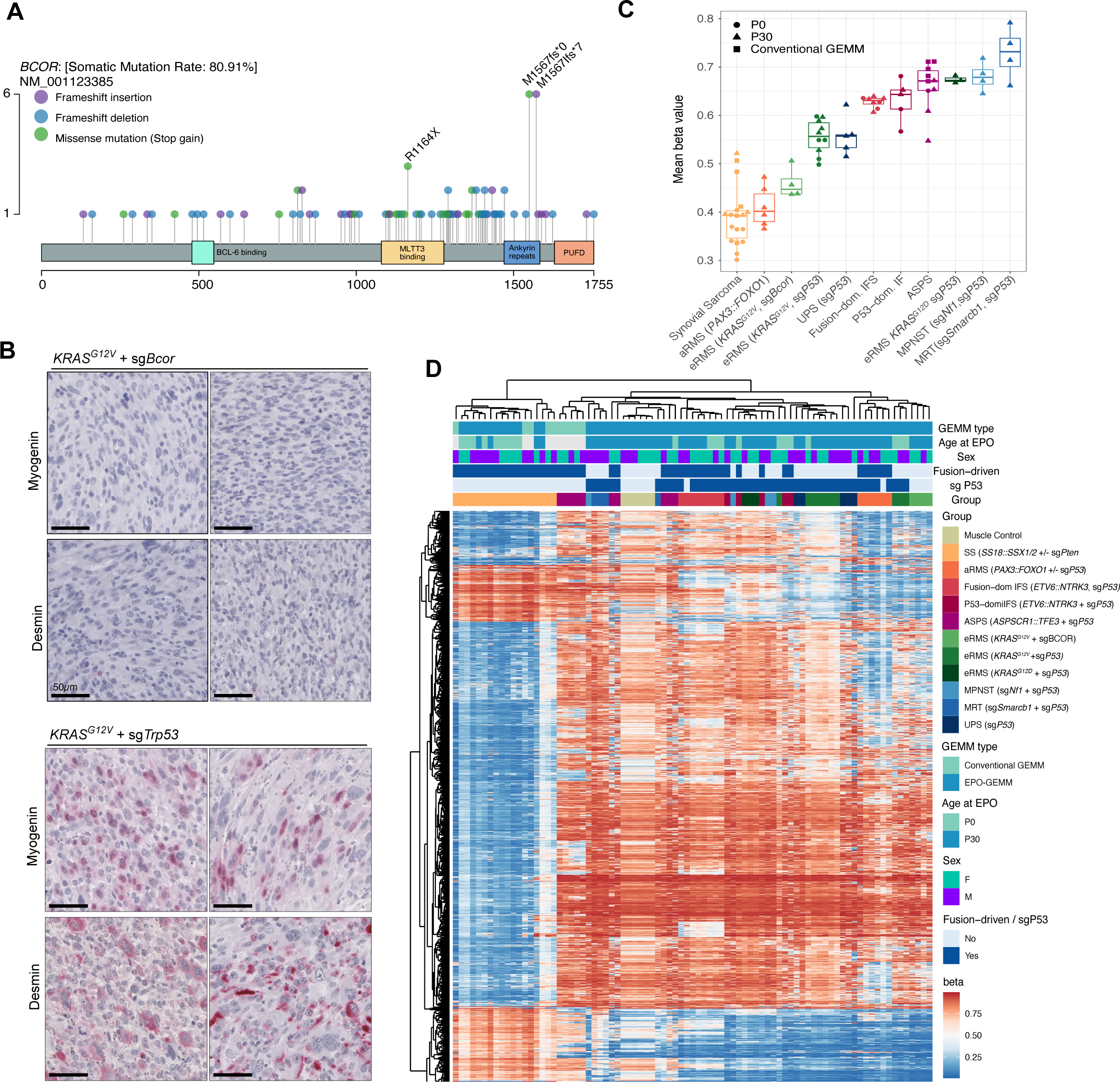
*BCOR* inactivation in RMS and DNA methylation patterns in mouse sarcomas A) BCOR mutations identified in human rhabdomyosarcoma, re-analysis of previously published data (Shern et al., 2021). B) Exemplary IHC histographs for mesenchymal and muscle-differentiation markers desmin and myogenin, across *KRAS^G12V^*/sg-*Bcor* and *KRAS^G12V^*/sg-*Trp53* tumors. C) Mean methylation beta values as boxplots ordered by median. D) Heatmap view of DNA methylation based on the top 10,000 differentially methylated CpG sites across the mouse sarcoma cohort. N ≥ 3 tumors per group.

**Figure S4.**
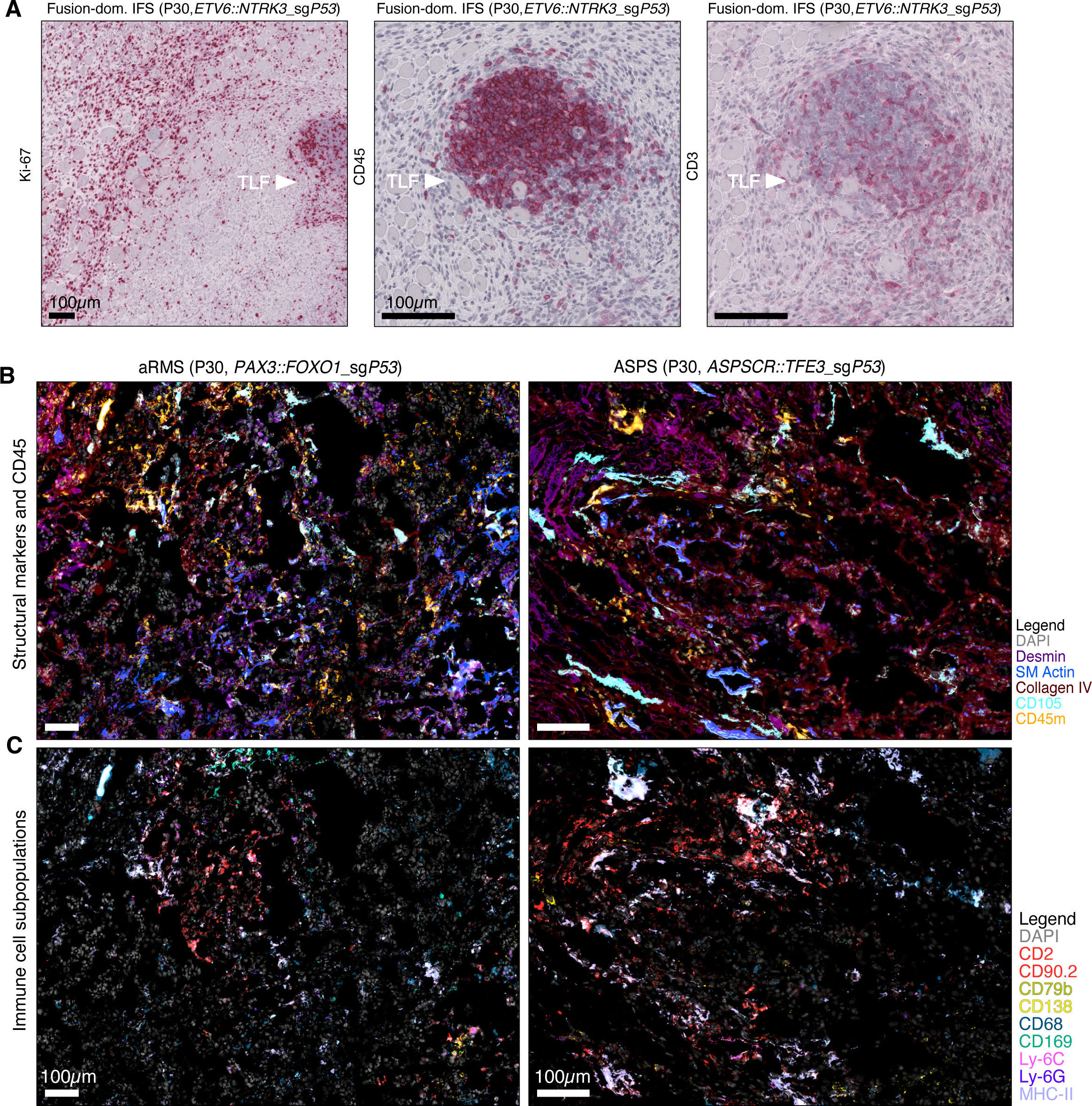
Immune infiltration phenotypes of mouse sarcomas **A)** Exemplary IHC histographs of immune infiltration in mouse sarcomas. TLF refers to tertiary lymphoid follicles, which were occasionally observed. Matching tumor regions in neighboring slides are depicted as H&E and IHC for Ki-67, CD45, CD3, demonstrating decreased proliferation rate of tumors cells in TLF proximity. **B-C)** Exemplary multiplex IF histographs of mouse sarcomas. Panels in **B** show structural markers, using DAPI for nuclear staining, mesenchymal marker desmin, cytoskeletal markers smooth muscle actin and collagen IV, endothelial marker CD105 and CD45 for immune cells. Panels in **C** illustrate immune cell subpopulations in matching regions using DAPI for nuclear staining, CD2 and CD90.2 for T cells, CD79 for B cells, CD138 for plasma cells, CD68 as a pan macrophage marker, CD169 for CD169-positive macrophages, Ly-6G and Ly-6C for granulocytic and monocytic myeloid-derived suppressor cells (g-/m-MDSCs) and MHCII. Scale bars=100µm.

**Figure S5.**
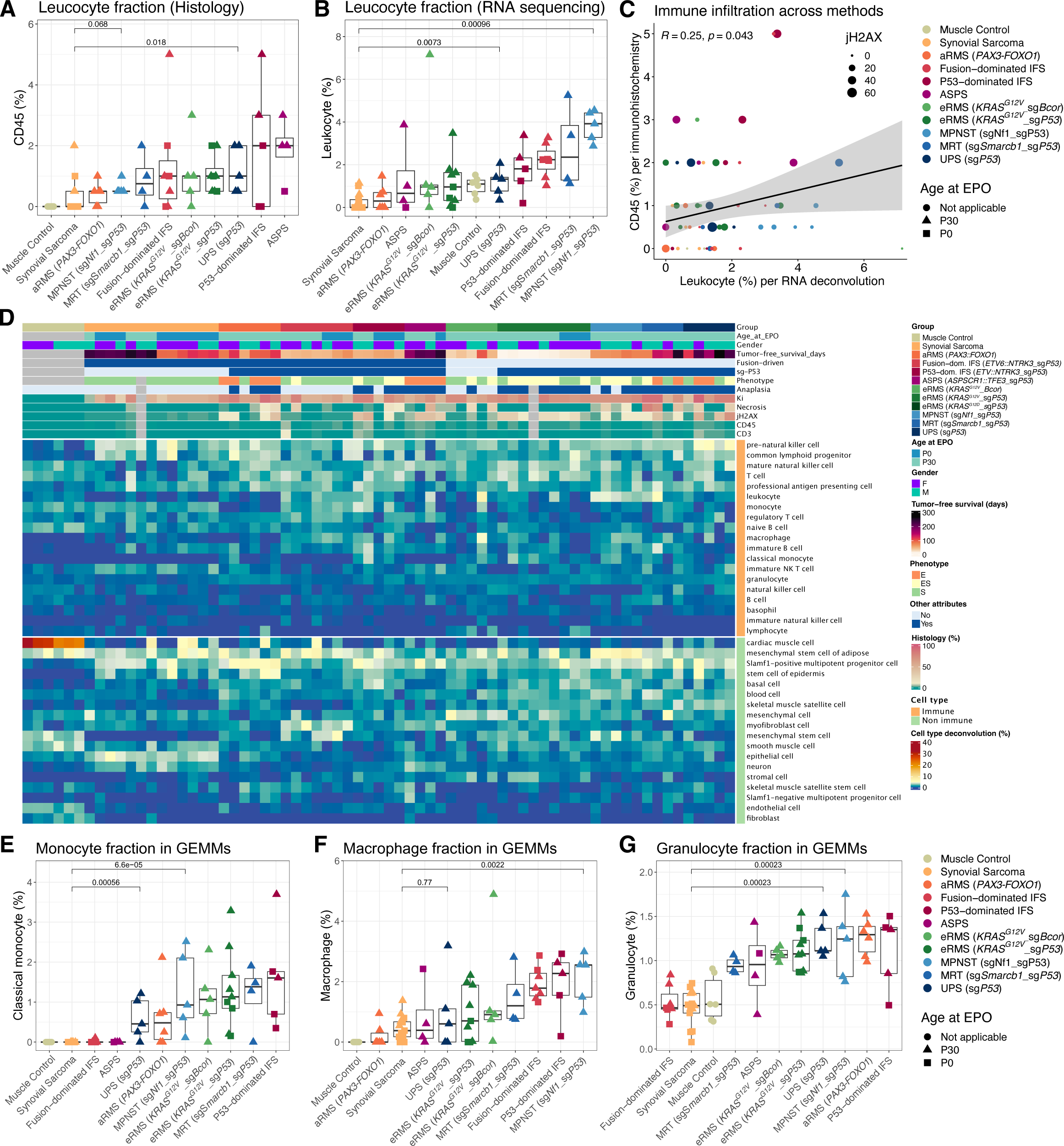
Immune cell deconvolution in mouse sarcomas A-C) Comparison of leukocyte quantification based on IHC (A) and cell type deconvolution (**B**) based on RNA sequencing using CibersortX and the publicly available single-cell RNA-seq Tabula muris data set as a reference matrix (Tabula Muris Consortium et al., 2018). Panel **C** shows a Pearson correlation between the two methods. **D)** Heatmap view of immune and non-immune cells using CibersortX and Tabula muris. **E-G)** Monocyte, macrophage and granulocyte fractions from panel **D** depicted as median-sorted boxplots. P-values of boxplots were determined by unpaired two-sided Wilcoxon tests and corrected for multiple testing by Bonferroni-Holm method. N ≥ 4 tumors per group.

**Figure S6.**
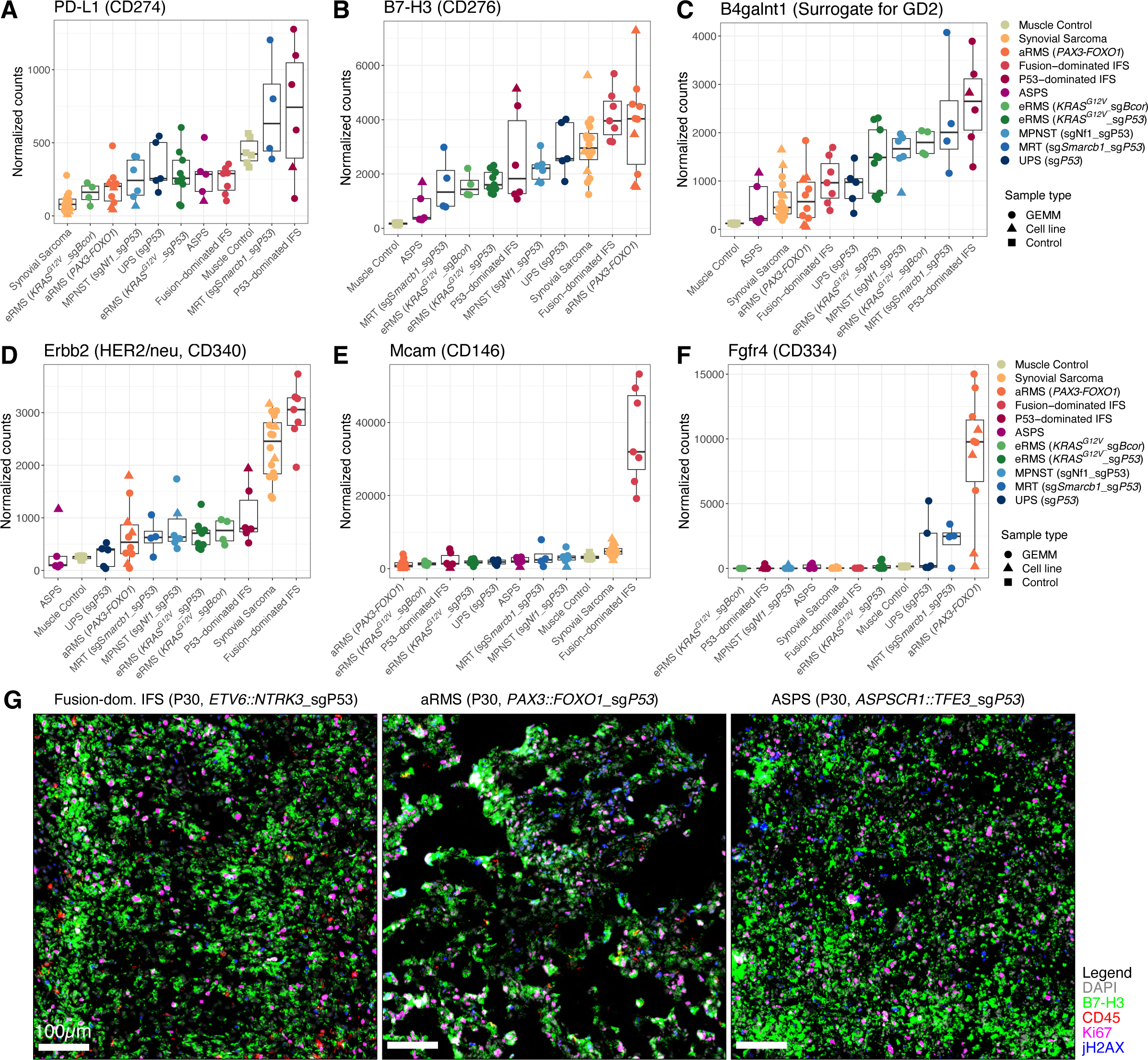
Expression of immunotherapy targets in mouse sarcomas A-F) Median-ordered boxplots, depicting normalized immunotherapy target gene expression across mouse sarcomas based on RNA-sequencing. The chosen pan-cancer and entity-specific targets, were previously identified in human cancer cohorts. For Ganglioside GD2, expression of producing enzyme B4galnt1 was plotted, previously established as a highly correlative marker gene (Sorokin et al., 2020). N ≥ 4 tumors per group. G) Exemplary multiplexed IF histographs depicting B7-H3 expression amidst DAPI for nuclear staining, CD45 for immune cells, jH2AX for DNA doubles strand breaks and proliferation markers Ki-67. Scale bars equal 100µm.

**Figure. S7.**
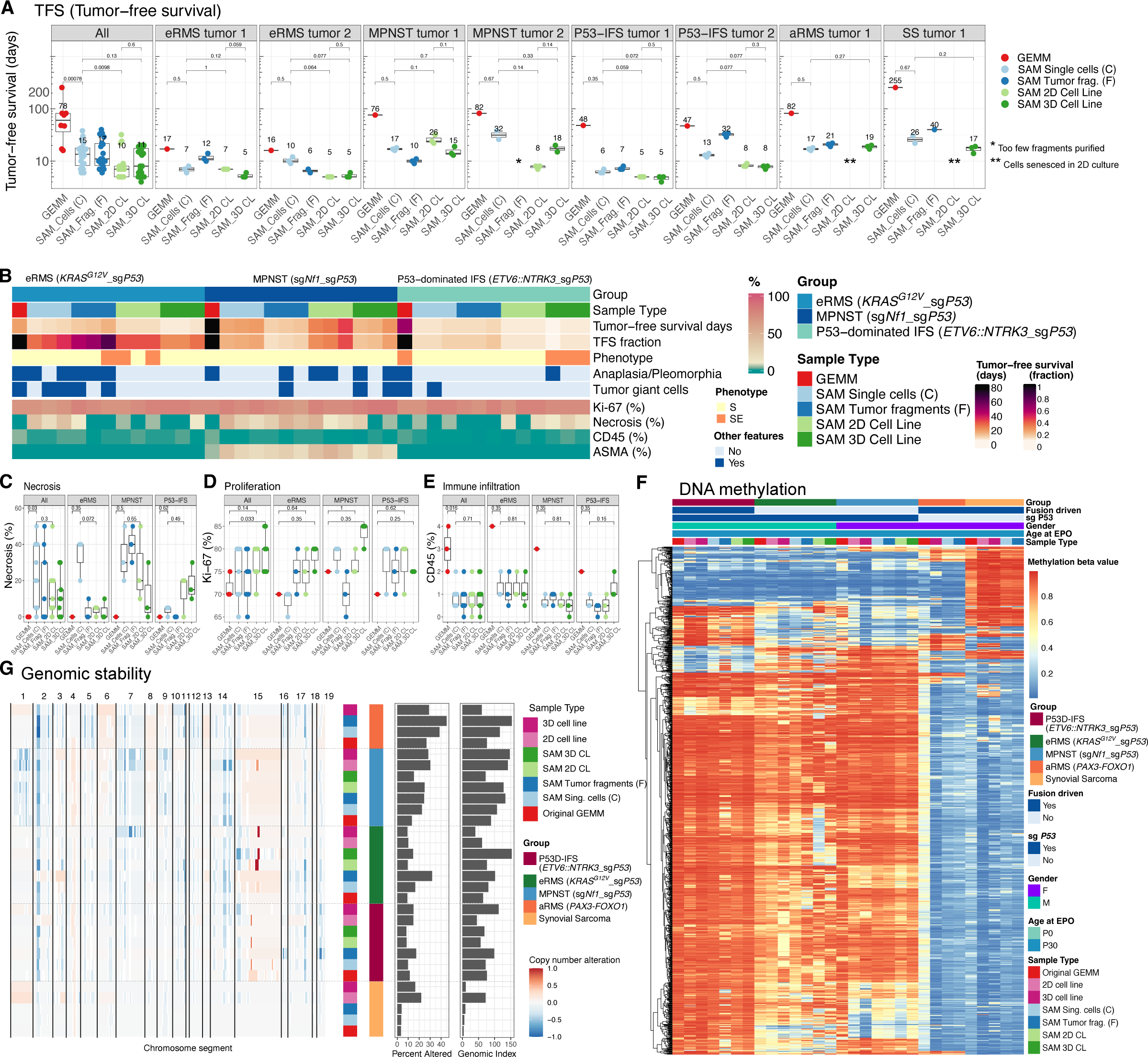
Syngeneic allograft models faithfully recapitulate GEMMs **A)** Tumor-free survival of SAMs compared to corresponding GEMMs. **B)** Heatmap view of mouse sarcomas of three different entities, systematically compared across original GEMM and different engraftment types, quantified for six morphological and IHC features by blinded expert pathology review. Asymmetric color scale for combined visualization of low (CD45) and high-scoring antigens. **C-E)** Histological features (Necrosis, Ki-67 and CD45 score) from panel **A** visualized as boxplots. P-values of boxplots were determined by unpaired two-sided Wilcoxon tests and corrected for multiple testing by Bonferroni-Holm method. N = 3 tumors per engraftment group. **F)** Heatmap view of DNA methylation based on the top 10,000 differentially methylated CpG sites comparing GEMMs and corresponding SAMs. N=1 tumor per group. **G)** Genomic stability in SAMs compared to GEMMs, visualized as copy number variations (CNV) profiles derived from DNA methylation data, condensed as Percent altered (≤/≥ .1) and Genomic Index. N=1 tumor per group.

**Figure S8.**
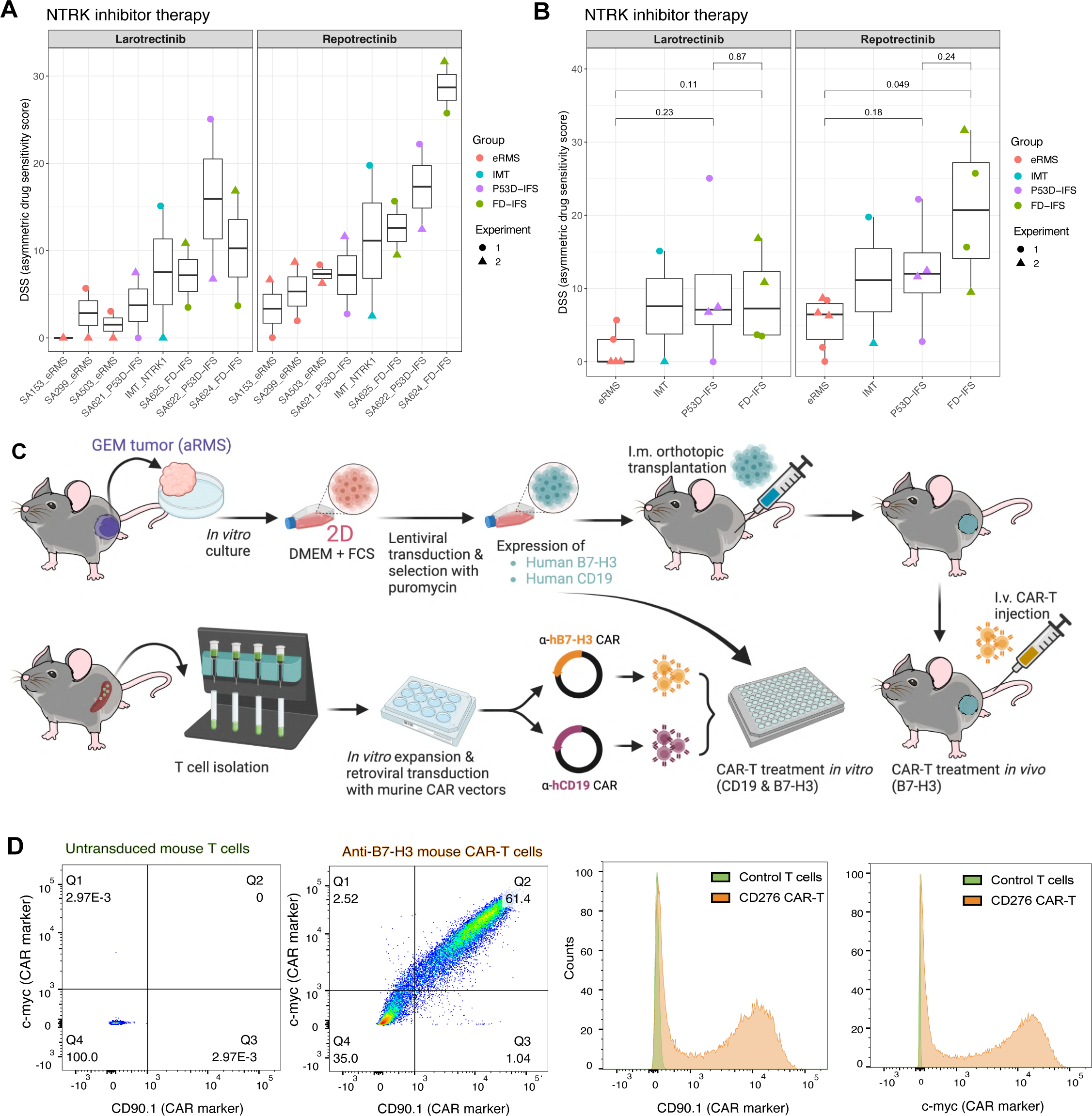
Translational model application A & B) Drug sensitivity scores from *in vitro* drug sensitivity testing for individual models (A) and grouped by entity (B). IMT refers to a human tumor cell line derived from an Inflammatory myofibroblastic tumor driven by *ETV6::NTRK3* used to clone the mouse transposon vectors for electroporation. P-values refer to unpaired two-sided t-tests, corrected for multiple testing using the Bonferroni-Holm method. C) Schematics of generation of murine CAR-T cells targeting human target antigens B7-H3 and CD19 used to treat a murine aRMS mouse tumor line lentivirally modified to express human B7-H3 and CD19. D) FACS analysis of mouse T lymphocytes (untransduced or transduced with B7-H3 CAR vector) stained for CAR markers CD90.1 and myc-tag and corresponding histograms.

